# A shared functional architecture for error-based and reinforcement-based motor learning in the human brain

**DOI:** 10.64898/2026.06.18.733155

**Authors:** Keanna Rowchan, Corson N. Areshenkoff, Ali Rezaei, Maryam Ansari Esfeh, Daniel J. Gale, J. Randall Flanagan, Jeffrey D. Wammes, Jason P. Gallivan

**Author notes:** Correspondence: Keanna Rowchan, Department of Psychology, Queen’s University. The two authors contributed equally to this work (Jeffrey D. Wammes, Jason P. Gallivan).

## Abstract

The motor system can flexibly learn from fundamentally different teaching signals, from directional errors to rewards. While computational and neurobiological models have long assigned these to distinct error-based learning (EL) and reinforcement-based learning (RL) processes, whether this separation holds at the level of whole-brain network architecture has not been directly tested. Here, we characterized whole-brain functional connectivity manifolds in the same participants as they performed separate EL and RL motor tasks — differing in both feedback type and motor structure — during two fMRI sessions. By jointly embedding covariance patterns from both tasks into a common low-dimensional neural space, we isolated task-general from task-specific learning-related network reconfigurations. We show that both forms of motor learning induce converging changes in the manifold structure of higher-order transmodal cortex, with comparatively limited task-specific modulation. Notably, this convergence extended to the cerebellum and basal ganglia, the canonical substrates of error-based and reward-based learning, which reconfigured comparably across both tasks. Against this shared backdrop, we found that the posterior medial cortex exhibited a unique functional geometry, selectively redistributing its connectivity across distinct brain circuits depending on feedback type and learning stage. We further demonstrate that individual differences in domain-general motor learning ability are associated with stage-dependent reconfigurations within limbic, default-mode, and attentional systems. These findings indicate that flexible motor adaptation, irrespective of the nature of the learning signal, is supported less by changes within task-specific sensorimotor circuits than by the dynamic reorganization of higher-order brain networks that coordinate them.

## INTRODUCTION

Learning mappings between motor commands and their sensory outcomes is fundamental to daily life, supporting the acquisition of new skills and adaptation to changes in the body or environment. Given the diversity of sensory signals that can drive this learning, it is not surprising that the motor system is widely believed to rely on multiple distinct learning processes, each preferentially tuned to different forms of sensory feedback (1–6). Two of the most extensively studied processes for mapping movement to feedback are error-based learning (EL) and reinforcement-based learning (RL). In EL, sensory prediction errors that encode directional discrepancy between actual versus expected sensory outcomes drive systematic adjustments of motor commands (6–8) — for instance, when a tennis player sees their serve land wide and corrects their arm trajectory on the next attempt. In RL, learning is guided instead by scalar reward signals that indicate the success or failure of an action, often without directional error information, requiring the motor system to explore possible actions and assign credit to those that maximize reward (6, 9, 10) — as when a child learning to ride a bicycle discovers, through trial and error, which combinations of balance and pedaling keep them upright. Because these two processes rely on such fundamentally different signals, it has long been assumed that they are driven by distinct, independent neural architectures.

In line with this assumption, computational and neurobiological models have traditionally proposed a strict division of labor between cortico-subcortical circuits. During EL, the cerebellum has been widely recognized as a key substrate for computing sensory prediction errors, operating through cortico-cerebellar pathways (5, 6, 11). During RL, in contrast, the basal ganglia has been associated with dopaminergic reward prediction errors that support action selection via cortico-striatal circuits (1, 4, 12–14). While this dual-systems framework has been highly influential, a strict anatomical dissociation between EL and RL circuits may provide an incomplete account of learning, overlooking the broader networks that support performance across both forms of learning (15, 16).

Consistent with the idea of shared mechanisms, a growing body of behavioral work has demonstrated that both EL and RL are shaped by higher-order cognitive processes that operate alongside, and often independently of, core error-correction mechanisms. In standard EL tasks, where individuals must adjust their movement trajectories to hit a target under distorted visual feedback, explicit re-aiming strategies account for a substantial portion of total adaptation and can be dissociated from implicit sensory recalibration (16–18). In RL tasks, where individuals must continuously shape their movement trajectories through reinforcement, the absence of directional error information makes trial-and-error search computationally expensive and the strategic narrowing of the action space accelerates learning (18–21). These observations present the possibility of a domain-general cognitive system — involving context monitoring, strategy formation, and performance evaluation — that supports flexible adaptation regardless of the specific feedback signal used to guide learning.

At the same time, accumulating evidence has complicated the classical neuroanatomical boundaries between EL and RL circuits at the subcortical level. The cerebellum, long considered central to the sensory prediction error processing that supports EL (5–7, 11, 22–24), has also been shown to encode reward prediction errors and exhibit reward-modulated activity (25–28). Similarly, the limbic cortices, which sit alongside the default mode network (DMN) at the apex of the cortical processing hierarchy (29, 30), have traditionally been associated with the reward valuation (31, 32) and memory processes (33, 34) that support RL. Yet recent work has also implicated these regions in error-based visuomotor adaptation at the level of individual differences (18), suggesting that their role in motor learning extends beyond classical RL. Together, these converging lines of evidence suggest that EL and RL may engage common patterns of large-scale brain network reorganization. This prediction, however, has not yet been tested directly within the same individuals.

Testing this prediction requires a framework to capture how learning reshapes distributed patterns of activity across cortical and subcortical systems simultaneously. Low-dimensional manifolds, derived from whole-brain functional connectivity (FC) patterns, provide such a framework (30, 35–37). By embedding the connectivity profile of every brain region into a common low-dimensional space, manifolds allow task-general and task-specific changes to be isolated within a single representational geometry — offering an analytic advantage over approaches that focus on predefined regions or pairwise connections. Applying this framework to motor learning has revealed that learning-related network reconfigurations extend well beyond sensorimotor cortex, engaging transmodal regions including the DMN (29–31), as well as limbic and cerebellar systems (27). However, these studies are limited by examining EL and RL in isolation and using separate samples, leaving unresolved whether these distinct forms of learning converge on shared neural trajectories within whole-brain connectivity space.

Here, we addressed this question using a within-subjects design where participants (N=37) performed an EL task (visuomotor adaptation) and RL task (reward-based trajectory shaping) during separate fMRI sessions. By embedding connectivity patterns from both tasks into a common low-dimensional manifold space, we isolated changes shared across tasks, identified deviations specific to EL or RL, and related individual differences in manifold reconfiguration to cross-task learning performance. We show that both forms of learning induce converging changes in manifold structure of higher-order transmodal cortex, and that this convergence extends to the cerebellum and basal ganglia — the canonical substrates of the classical EL/RL dichotomy. Against this task-general backdrop, the posterior medial cortex occupies a unique position, as the only region reflecting both task-dependent and task-general dynamics. We further demonstrate that individual differences in domain-general learning ability are associated not with changes in sensorimotor connectivity, but with learning-related reconfigurations of limbic and attentional networks. Our findings establish a direct link between higher-order brain network dynamics and individual differences in motor learning performance, and identify specific connectivity architectures in which the brain supports flexible adaptation across distinct forms of sensory feedback.

## RESULTS

### A shared dimension of learning performance across EL and RL

Participants (N=37) performed an EL and RL task during two fMRI sessions one week apart. Both tasks required participants to move a cursor, representing their finger position, to intercept a target using an MRI-compatible touchpad, subject to different forms of visual feedback (for an overview of task designs and procedures, refer to our *Methods and Materials*).

In the EL task, participants performed a classic visuomotor rotation task (7, 16, 17), making center-out movements along the touchpad with their right index finger toward one of eight targets arranged on a circular ring, while being provided cursor feedback about their index finger’s position (see Fig. 1A). After a baseline block (64 trials), participants completed a learning block (160 trials) where cursor feedback was rotated clockwise by 45° relative to the finger’s actual trajectory. Over trials, participants learned to adapt to this visuomotor rotation, wherein they adjusted their trajectory to compensate for the 45° perturbation (see Fig. 1B; Fig. 1C for example participant).

**Figure 1.**
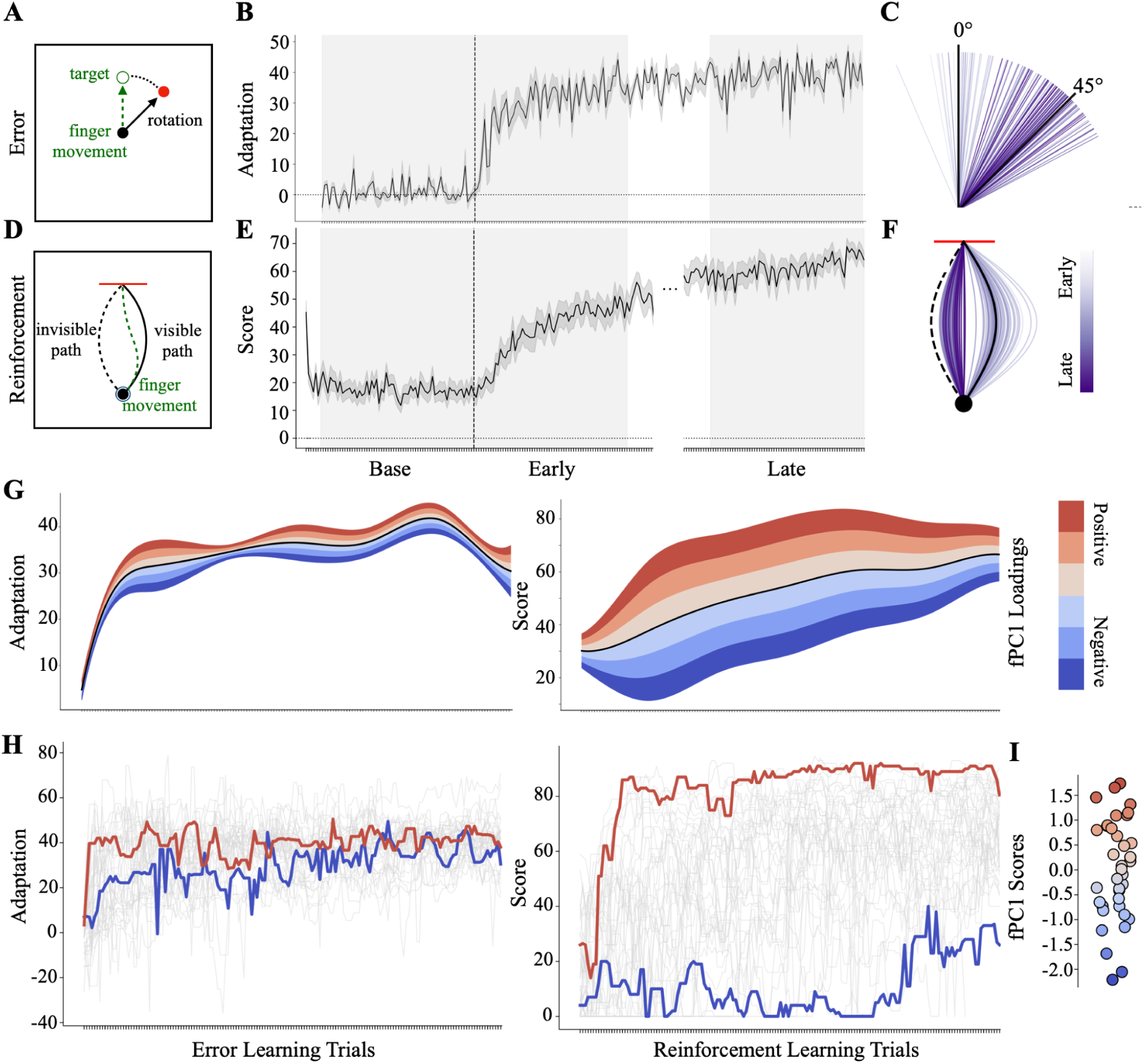
A task-general dimension of learning ability describes inter-individual variability across error-based and reinforcement-based motor learning. (A) Trial structure of the EL task. During learning, cursor feedback was rotated 45° relative to hand movement direction, requiring participants to adapt their reach direction. The green arrow indicates the participant’s intended hand movement direction, while the black arrow represents the rotated cursor feedback (45° rotation). The white circle with green outline denotes target location whereas the red circle denotes endpoint of the rotated cursor position. (B) Group adaptation curves in the EL task. The black line denotes mean adaptation (45° − reach error); shaded regions denote standard error of the mean (SEM). For the neural analyses, the learning block was divided into 64-trial Early and Late epochs. (C) Adaptation across trials for a single participant, with color intensity reflecting progression from Early- (light purple) to Late-learning (dark purple). (D) Trial structure of the RL task. Participants traced a visible curved path (black line) and received scalar feedback (0–100) based on tracing accuracy relative to an unseen mirror-reversed target path (dashed black line). The green dashed line reflects participants’ hand movements from the start position (filled black circle) to the target line (red). (E) Group-averaged learning curve in the RL task. (F) Reach trajectories for a single participant in the RL task. (G) Functional PCA (fPCA) applied to individual learning curves across both tasks. A single component (fPC1) was found to capture 61.2% of the variance across tasks, reflecting a shared dimension of learning ability (i.e., ʻcross-task learning score’). Shaded bands show the effect of positive (red) and negative (blue) cross-task learning scores relative to the group mean (black trace). (H) Learning trajectories from two participants who exhibited the highest (red) and lowest (blue) cross-task learning scores, shown across both tasks. (I) fPC1 loadings for individual participants.

For the RL task, we modified a classic motor paradigm (20) in which participants used their right index finger to trace a rightward-curved path on the touchpad without visual cursor feedback (see Fig. 1D). Following a baseline block (70 trials) where no feedback was provided, participants performed a learning block (200 trials) in which they received scalar reward feedback (0–100) reflecting tracing accuracy. Unbeknownst to participants, however, feedback was computed based on their adherence to a mirror-reversed version of the target path, instead of the actual target path. As cursor feedback was unavailable, learning relied solely on this scalar reinforcement feedback. At the group-level, scores increased across trials, reflecting learning (see Fig. 1E; Fig. 1F for example participant).

Although group-averaged performance confirmed successful learning in both tasks, mean learning curves can obscure substantial inter-subject variability in both the rate and extent of learning (17, 18, 41). To isolate dominant sources of this variability, we applied functional principal component analysis (fPCA; <u>42</u>; see Methods) to subjects’ individual learning curves across both of the tasks (see Fig. 1G). We found that a single functional principal component (fPC1) accounted for the majority (61.2%) of the variance in learning trajectories, capturing differences in overall learning performance across individuals. Accordingly, participants’ scores on fPC1 reliably distinguished good from poor learners across tasks (see Fig. 1H–I; Supplementary Fig. 1-2 for robustness of fPC measures), and we hereafter refer to this measure as participants’ *cross-task learning score*.

To assess how each task contributed to this shared dimension of learning, we performed fPCA separately on EL and RL learning curves and correlated, across subjects, task-specific scores with the cross-task learning score (see Supplementary Fig. 2). RL contributed the majority of the inter-individual variance (r = 0.998 with the combined score), consistent with greater behavioral variability due to the absence of directional error information in this task (see Fig. 1G, Right, relative to Left). EL showed a weaker but significant positive association (r = 0.327, p = 0.048), indicating that the cross-task learning score captures shared learning variability across tasks. We therefore used cross-task learning scores as the primary behavioural index of learning performance across tasks in all subsequent neural analyses.

This shared behavioral dimension is consistent with prior work implicating common cognitive processes across motor learning paradigms (15, 19, 43–45), and raises the possibility that domain-general learning ability may be supported by higher-order brain networks that operate above task-specific sensorimotor circuits. We return to this question later in our neural analyses, after first characterizing the whole-brain manifold architecture of learning across both tasks.

### Whole-brain manifold captures canonical functional organization

To characterize learning-related changes in whole-brain connectivity, we estimated FC manifolds from cortical (Schaefer 400 parcellation; <u>46</u>), subcortical (Tian 32-region atlas; <u>47</u>), and cerebellar (Nettekoven 32-region atlas; <u>48</u>) timeseries data across one resting-state and six equal-length epochs across the EL and RL tasks (Baseline, Early-, and Late-Learning windows for each task; 198 imaging volumes each; Fig. 2A-B). Prior work has demonstrated that the majority of variance in whole-brain FC patterns reflects static, trait-like individual differences rather than state- or task-related modulation (49, 50), and these subject-level features can mask subtle connectivity changes associated with different task states (44, 51). To address this, we applied Riemannian manifold centering (<u>51–53</u>; see Methods), which removes subject-level variance in FC patterns. The effectiveness of this procedure is illustrated in Fig. 2D: prior to centering, FC matrices clustered mainly according to subject identity; following centering, this subject-level clustering was eliminated, exposing task- and epoch-related variation. We then constructed low-dimensional manifolds via PCA for each participant and epoch, and aligned all manifolds to a common resting-state template using Procrustes transformation (see Fig. 2A–C). This template was constructed from resting-state FC data acquired from our same participants (see <u>37–40</u> for similar approaches).

**Figure 2.**
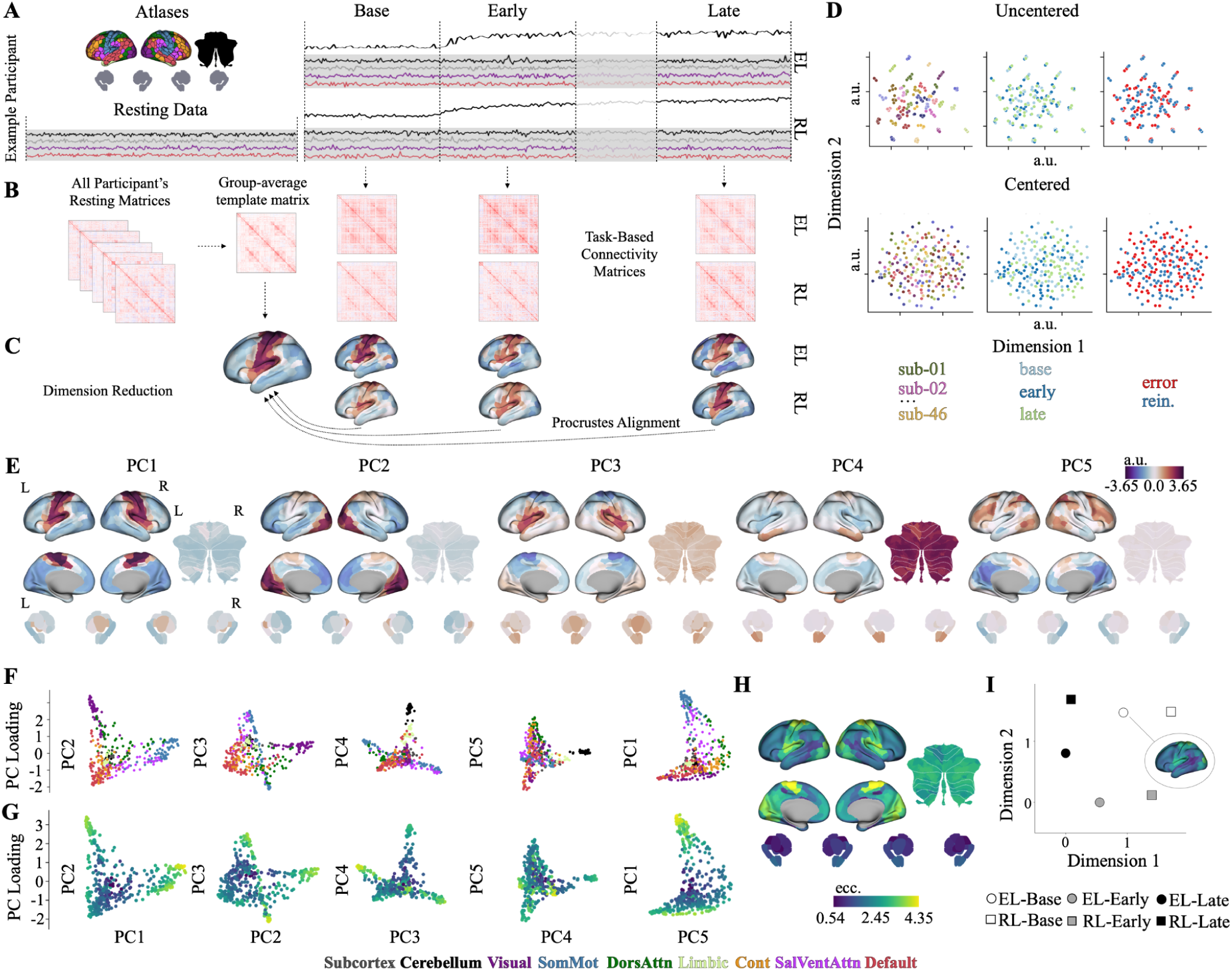
Whole-brain connectivity manifolds capture canonical axes of functional organization, and learning-related neural states cluster by epoch rather than by task. (A) For each participant, task, and epoch, regional BOLD timeseries were extracted from cortical (46), subcortical (47), and cerebellar (48) regions. (B) FC matrices were estimated for each participant and epoch. For resting-state data — collected from the same participants — FC matrices were averaged to create a common, group-average template. (C) PCA was applied to each matrix to construct low-dimensional manifolds. All manifolds were aligned to the group-average resting-state template using Procrustes alignment. (D) UMAP visualization (57) of FC matrices before (top) and after (bottom) Riemannian manifold centering, colored by subject identity (left), epoch (middle) and task (right). Prior to centering, matrices clustered by participant; following centering, this subject-level structure was eliminated. (E) Whole-brain loadings for the top five PCs of the resting-state template manifold, projected onto surface space. The top five PCs accounted for the majority (65.4%) of variance. (F) Each brain region depicted as a point in the five-dimensional manifold space (projected into 2D for visualization), colored by Yeo 7-network assignment (46), with subcortical (grey) and cerebellar (black) regions. (G) Regional manifold eccentricity (i.e., Euclidean distance from the manifold centroid) visualized within low-dimensional manifold space. Higher eccentricity (yellow) indicates greater functional segregation; lower eccentricity (dark blue) indicates greater functional integration (see Supplementary Fig 4). (H) Regional eccentricity values projected onto surface space. (I) UMAP visualization of whole-brain eccentricity profiles across subjects, tasks, and epochs. Each point represents the group-mean eccentricity pattern for a given task epoch. Note eccentricity patterns tend to cluster by epoch rather than by task, providing an initial indication that learning-related manifold reorganization might be shared across tasks.

The top five PCs of the resting-state template manifold capture the canonical axes of large-scale brain organization, accounting for 65.4% of the variance in FC (see Fig. 2E–F). We retained these five components based on a clear inflection point in the variance-explained curve (see Supplementary Fig. 3), beyond which additional components contributed diminishing explanatory power (a standard criterion for dimensionality selection in manifold analyses; <u>36, 54, 55</u>). PC1 separated unimodal sensorimotor regions from visual cortex and transmodal cortex, including the DMN; PC2 distinguished visual cortex from transmodal regions; and PC3–5 captured finer distinctions among attention, control, DMN, and cerebellar-hippocampal systems. To reduce this multivariate embedding to a single scalar index, using established approaches (38, 39, 56), we computed ‘manifold eccentricity’ — the Euclidean distance of each region from the manifold centroid — for each participant, region, and epoch (see Fig. 2G–H). Regions with higher eccentricity occupy the periphery of the manifold and exhibit more functionally segregated connectivity profiles, whereas regions with lower eccentricity are closer to the center and exhibit greater functional integration (see Supplementary Fig. 4). Notably, when we projected whole-brain eccentricity profiles for each learning task epoch into a Uniform Manifold Approximation and Projection (UMAP; <u>57</u>) space, we found that eccentricity patterns appeared to primarily cluster by learning epoch rather than by task (see Fig. 2I), providing an initial indication that learning-related manifold reorganization might be shared across tasks.

### Learning-related manifold changes in cortex are shared across EL and RL

Consistent with the qualitative pattern from the UMAP projection, we found that learning-related changes in manifold eccentricity were broadly shared across tasks. A region-wise rmANOVA with Task (EL, RL) and Epoch (Base, Early, Late) as within-subject factors revealed widespread main effects of Epoch across cortex (252 regions), with only 11 cortical regions exhibiting a main effect of Task (FDR-corrected, q < 0.05; see Fig. 3A–B). Subcortical and cerebellar regions also showed robust Epoch effects, which we address in the following section. Critically, no regions showed a significant interaction, indicating that the pattern of changes in manifold eccentricity across learning epochs was similar across tasks (see Supplementary Fig. 5 for full distributions of F-values across main effects and the interaction). Follow-up paired t-tests (FDR-corrected) on the Task effect revealed that these regions — concentrated in and around the posterior cingulate cortex (PCC) and precuneus — exhibited significantly lower eccentricity during EL relative to RL (see Fig. 3C). The anatomical specificity of this effect foreshadows a distinctive role for the posterior medial cortex (PMC) which we examine further in a separate subsection below.

**Figure 3.**
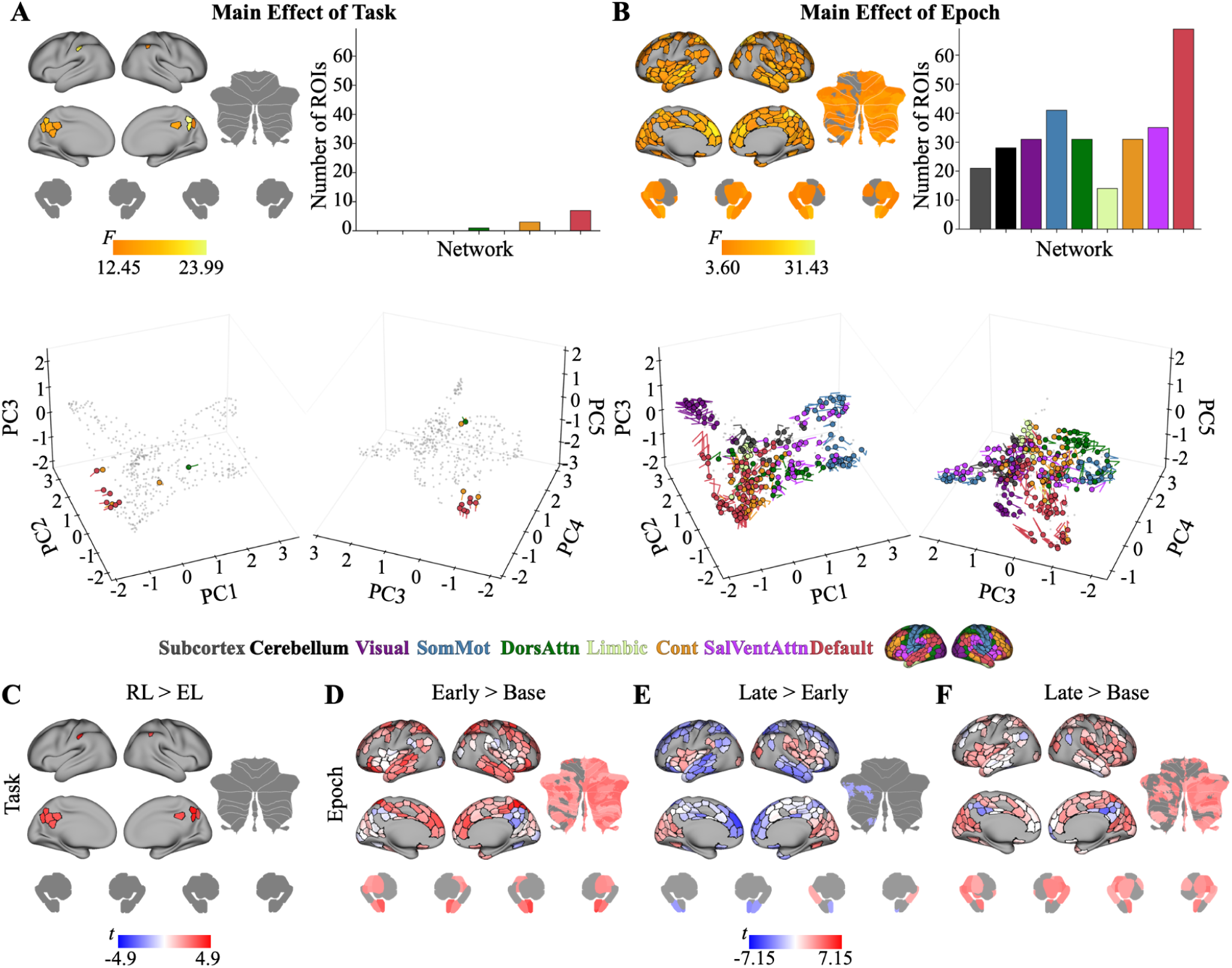
Learning-related manifold changes are shared across EL and RL, with task-specific effects confined to the posterior medial cortex. (A) Brain regions exhibiting a significant main effect of Task (EL vs. RL) and (B) a significant main effect of Epoch (Base, Early, Late) on manifold eccentricity, projected onto surface space. Epoch effects (301 regions spanning cortex, subcortex, and cerebellum) outnumber Task effects (11 regions, all cortical); no regions exhibited a significant interaction. Bar plots (at right) show the number of significant regions by Yeo 7-network assignment (46), including cerebellar (black) and subcortical (grey) regions. Below, Temporal trajectories of significant regions are shown in 3D manifold space, colored according to its functional network assignment. For the main effect of Task (panel A), coloured circles indicate each region’s initial position during EL, with the traces showing displacements during RL. For the main effect of Epoch (panel B), coloured circles indicate each region’s initial position during Baseline, and traces show their unfolding displacements during Early and Late-learning. (C) Pairwise comparison between the two tasks, collapsed across epochs. Regions in the PMC exhibited significantly lower eccentricity during EL, suggesting increased functional integration during error-based learning. (D) Early>Base contrast, collapsed across tasks. Widespread increases in eccentricity (red) indicate functional segregation across regions at the onset of learning, with the PMC region as a notable exception, tending to show decreased eccentricity (integration) instead (blue). (E) Late>Early contrast. Higher-order transmodal regions (limbic, control, DMN) showed decreased eccentricity, whereas visual and SalVentAttn regions showed increases — indicating a redistribution of integration toward transmodal networks as learning progressed. (F) Late>Base contrast. The PMC exhibited sustained decreases in eccentricity across learning. Notably, the PMC was the only region to exhibit both task and epoch sensitivity (panels A and B). All results FDR-corrected (q < 0.05).

The predominance of Epoch over Task effects (as illustrated by contrasting Fig. 3A and 3B; Supplementary Fig. 5), and absence of interactions, indicates that motor learning is accompanied by a broad, task-general reorganization of FC, with comparatively limited modulation specific to either feedback type. This convergence is notable given that the two tasks differed not only in feedback type (i.e., directional errors in EL versus scalar rewards in RL) but in their fundamental motor structure (i.e., discrete target-directed reaching in EL versus continuous path tracing in RL). This suggests that the shared epoch effects reflect common properties of the learning process itself rather than common motor demands. We next characterized these shared Epoch effects through pairwise comparisons aggregated across tasks, focusing on cortical dynamics here and returning to subcortical and cerebellar patterns in the following subsection.

At the transition from Baseline to the onset of learning (Early>Base; see Fig. 3D), we observed significant increases in manifold eccentricity — indicative of greater functional segregation — across widespread cortical regions, including somatomotor cortex, the dorsal attention network (DAN), and higher-order DMN and limbic networks in particular. In contrast, the PMC tended to exhibit significant decreases in eccentricity, indicating increased functional integration.

As learning progressed from Early- to Late-learning (see Fig. 3E), we observed a reversal of this pattern. Regions within the visual cortex and Salience/Ventral Attention (SalVentAttn) network showed increased eccentricity, whereas higher-order regions within the limbic, control, and DMN networks that had previously increased in eccentricity now displayed significant decreases, suggesting a redistribution of integration toward transmodal networks as performance stabilizes in Late-learning.

Finally, when we compared Base- to Late-learning directly (see Fig. 3F), we found that a majority of cortical regions exhibiting early increases in eccentricity did not fully return to Baseline levels during Late-learning. That is, despite the partial reversals observed from Early- to Late-learning, these cortical regions remained more segregated than their pre-learning state (red regions in Fig. 3F). However, select regions within the DMN — including the PMC and angular gyrus — instead showed sustained decreases in eccentricity from Base through Late-learning, reflecting a durable shift toward greater integration that persisted across the full timecourse of learning. These persistent patterns are particularly notable in the context of EL, where behavioral errors during Late-learning had returned to near-baseline levels (with participants achieving near-complete, ∼45° adaptation). This means the persistent connectivity changes cannot be attributed to ongoing error processing, and instead likely reflect the stable network architecture of the adapted motor state. That these same Base-to-Late changes were shared across EL and RL despite the complete absence of directional error signals in RL reinforces this interpretation: the late-learning manifold configuration likely captures a feedback-independent signature of motor learning, not an artifact of any particular error signal.

### Cerebellar and subcortical reconfigurations parallel cortical dynamics across both tasks

As described in the introduction, classical models of motor learning assign error-based and reinforcement-based processes to distinct subcortical substrates: the cerebellum for EL via sensory prediction errors (5, 6, 11), and the basal ganglia for RL via dopaminergic reward prediction errors (1, 4, 12–14). Our manifold results allow a direct test of this prediction at the whole-brain level, as both structures were included in our whole-brain analyses. Of the 28 cerebellar and 21 subcortical regions showing significant Epoch effects, none showed a significant main effect of Task or a significant Task × Epoch interaction (FDR-corrected, q < 0.05; compare Fig. 3A vs. 3B). Thus, at the level of whole-brain manifold organization, the cerebellum and basal ganglia reconfigured comparably across tasks, and did so along shared trajectories across learning stages. This pattern runs counter to the anatomical dissociation predicted by dual-systems models.

These trajectories were also non-monotonic, mirroring the cortical dynamics described above. From Base- to Early-learning, nearly the entire cerebellum expanded along the manifold, alongside eccentricity increases in the hippocampus, amygdala and thalamus — indicating widespread functional segregation at the onset of learning in both tasks (Fig. 3D). From Early- to Late-learning, this pattern partially reversed, with the amygdala and left cognitive/association cerebellar regions exhibiting manifold contraction (Fig. 3E). Finally, our Base-to-Late contrast revealed that large swaths of the cerebellum — particularly in association/cognitive zones — remained in a persistently segregated state, along with the thalamus, amygdala, and hippocampus. This mirrors the persistent manifold expansion observed in many cortical regions (Fig. 3F) and indicates that subcortical FC changes accompanying motor learning are not transient reactions to novelty but durable reconfigurations that outlast the initial acquisition phase.

We next turn to the set of regions in the brain that distinguished EL from RL — the posterior medial cortex (PMC) — and examine how it selectively couples with functional subdivisions within these same subcortical structures as a function of the learning context.

### The PMC exhibits both task-dependent and learning stage-dependent connectivity

The PMC (encompassing five PCC and one precuneus region) was the only cortical area to exhibit significant main effects of both Task and Epoch in our rmANOVA (see Fig. 3A–B). To understand what drives this unique dual sensitivity, we used these PMC regions as seed regions to examine how their connectivity profiles change across tasks and learning stages. Because manifold eccentricity reflects a multivariate shift in a region’s *whole-brain* connectivity profile, no single pairwise connection constitutes a direct test of what drives an eccentricity change. We therefore present unthresholded seed-based connectivity maps throughout, which capture the full spatial distribution of connectivity changes underlying the observed eccentricity effects.

During RL, the PMC showed stronger connectivity with other DMN regions, the posterior hippocampus, and right (ipsilateral to the hand) DMN-associated cerebellar regions (red regions in Fig. 4E; see Fig. 4F for these findings summarized at the network-level). By contrast, during EL, the PMC exhibited greater coupling with regions across fronto-parietal and sensorimotor cortex, the caudate and putamen, and right (ipsilateral to the hand) action-related zones of the cerebellum (blue regions in Fig. 4E; see also Fig. 4F). Note that these regions did not show either a global increase or decrease in connectivity in one task relative to the other — a pattern of quantitative changes that could, in principle, have explained the eccentricity difference alone. Instead, the PMC exhibited a qualitative redistribution of connectivity across distinct brain regions depending on the task: coupling with sensorimotor cortex and the dorsal striatum during EL, while coupling with DMN regions and the hippocampus during RL. This selectivity was especially pronounced within the cerebellum. Although cerebellar regions as a whole did not distinguish tasks at the manifold level (see preceding subsection), the PMC differentially engaged distinct functional subdivisions of the cerebellum depending on task: action-related motor zones during EL, and association/cognitive zones in RL. The same selectivity was evident within the striatum, where the PMC coupled preferentially with dorsal striatal subdivisions during EL. These patterns indicate that the PMC engages functionally specific subcircuits within the cerebellum and striatum as a function of the sensory signal being used to guide learning.

**Figure 4.**
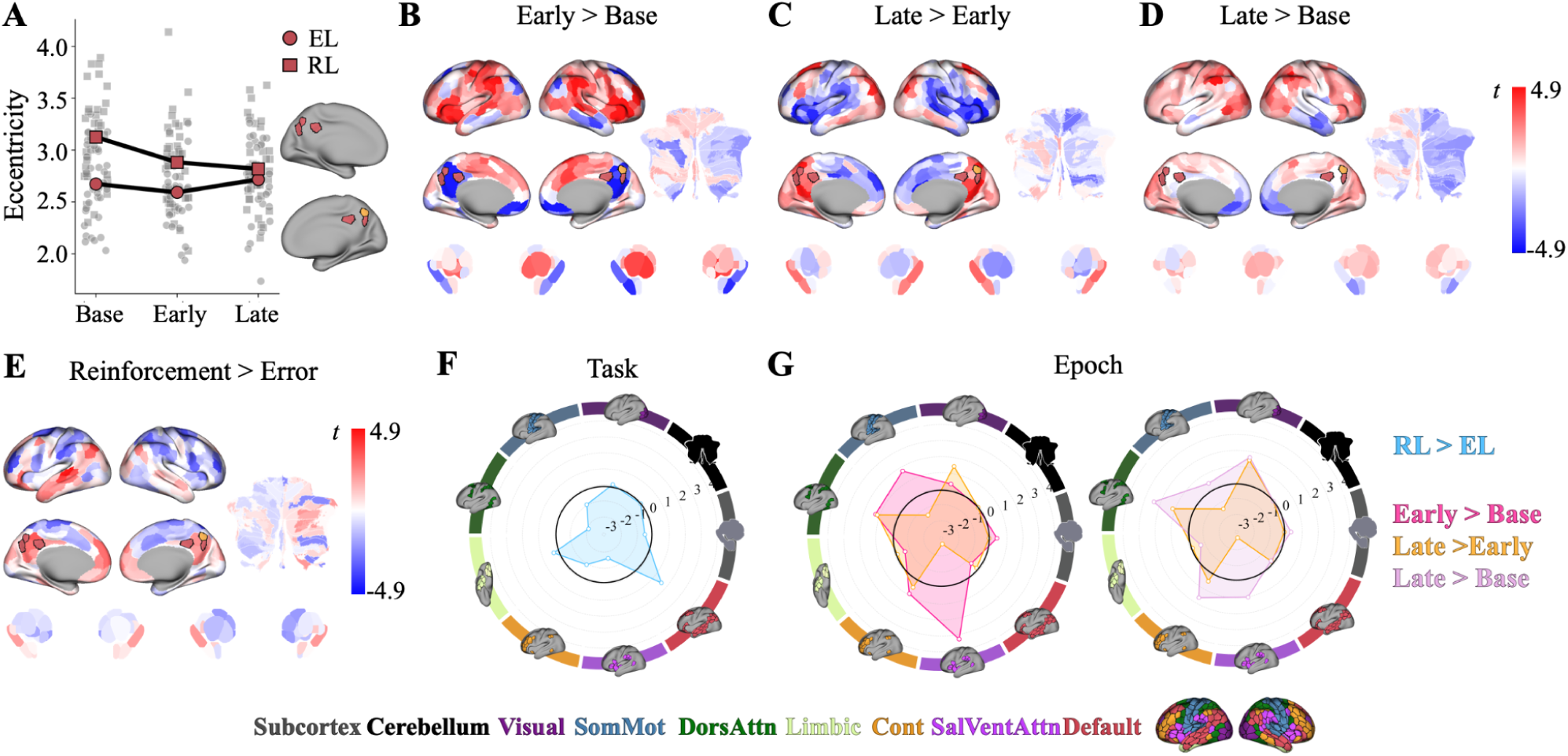
The PMC reconfigures its connectivity as a function of both task type and learning stage. (A) Mean PMC eccentricity averaged across regions for each subject, shown across tasks (EL, RL) and learning epochs (Base, Early, Late), with individual participants in grey. Six PMC regions (five PCC and one precuneus, projected onto surface space at right) were the only regions to exhibit both significant Task and Epoch effects. Note that PMC regions are color-coded according to their Yeo 7-network assignment (46) (see legend at bottom). (B–D) Seed-based connectivity changes for PMC regions across epoch contrasts: Early>Base (B), Late>Early (C), and Late>Base (D). Red indicates increased connectivity with PMC seed regions; blue indicates decreased connectivity. From Base- to Early-learning, the PMC increased connectivity with somatomotor and attentional networks. From Early- to Late-learning, this shifted toward greater connectivity with visual and transmodal regions. From Base- to Late-learning, the PMC showed sustained increases in connectivity with attentional and control networks. (E) Seed-based connectivity differences between tasks (RL>EL), collapsed across epochs. During RL, the PMC showed stronger connectivity with DMN regions, the hippocampus, and right DMN-associated cerebellar regions. During EL, the PMC coupled more strongly with fronto-parietal cortex, the caudate and putamen, and right action-related cerebellar regions. Significant regions from panel A are outlined in black. (F–G) Polar plots summarizing seed-based connectivity changes at the network-level for Task (F) and Epoch (G) contrasts. Note that, in panel G, the Late > Early contrast is maintained across both polar plots to allow for direct comparison. Network assignments follow the Yeo 7-network parcellation, with subcortical and cerebellar regions in gray and black, respectively.

The PMC’s connectivity also reconfigured dynamically across learning stages (see Fig. 4B–D, G). From Base- to Early-learning, PMC regions strongly decreased their connectivity with the hippocampus and other DMN regions — the network to which most of these parcels canonically belong — while simultaneously increasing FC with somatomotor and attentional networks, as well as the caudate and putamen. This pattern suggests the PMC partially disengages from its DMN affiliation at the onset of learning to interface with task-relevant sensorimotor systems. From Early- to Late-learning, this pattern largely reversed: PMC regions decreased connectivity with these same task-specific systems and increased communication with visual and higher-order transmodal regions, including the hippocampus, consistent with a gradual return toward broader within-network integration as performance stabilized. From Base- to Late-learning, the PMC showed sustained increases in connectivity with attentional and control networks, reflecting a longer-term reorganization that persisted even as behavioral performance — particularly in EL, where errors had returned to near-baseline levels — appeared outwardly stable. This profile indicates that the PMC dynamically redistributes its network coupling as a function of both learning type and stage, and that several of these changes persist after learning has largely plateaued.

### Limbic and attentional network reconfigurations are associated with cross-task learning

Having established the group-level manifold architecture of learning (in Fig. 3), we next asked whether individual differences in manifold reconfiguration were associated with cross-task learning performance. To this end, we correlated participants’ cross-task learning scores with regional changes in eccentricity across our key epoch contrasts (Early>Base, Late>Early, Late>Base; see Fig. 5A–C). Although no single region survived FDR correction, the resulting maps displayed notable spatial contiguity, suggesting that behaviourally relevant effects may instead be organized at the network (rather than single-region) level. We therefore aggregated regions within the Yeo 7-network parcellation (46) and evaluated network-level associations using spin permutation testing (1,000 rotations), which preserves cortical spatial autocorrelation (<u>58</u>; see Methods; Fig. 5D–F).

**Figure 5.**
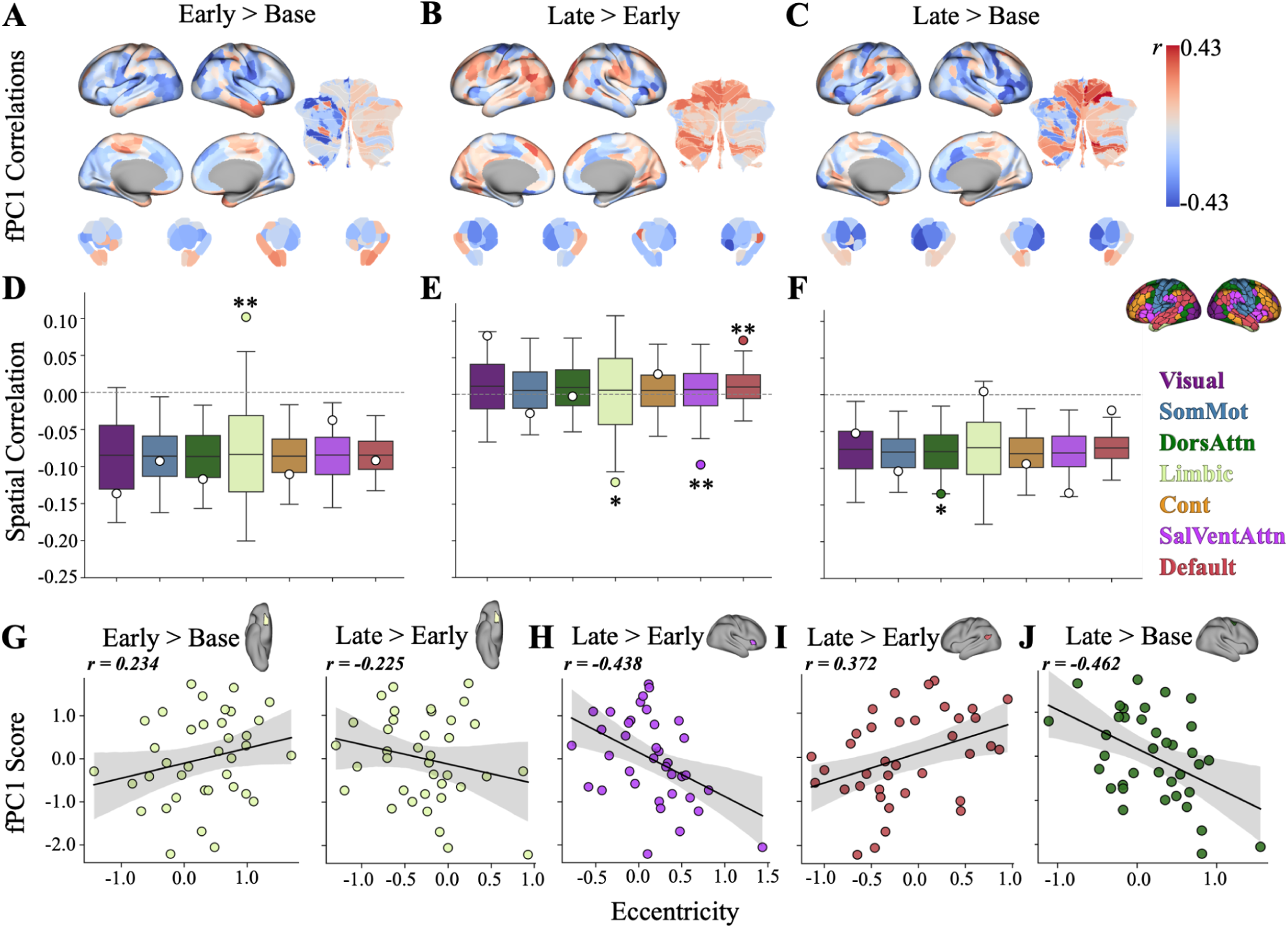
Individual differences in cross-task learning performance are associated with stage-dependent reconfigurations within limbic and attentional networks. (A–C) Whole-brain maps showing correlations between participants’ cross-task learning scores and changes in regional manifold eccentricity for each epoch contrast: Early > Base (A), Late > Early (B), Late > Base (C). (D–F) Network-level statistical tests using spatial autocorrelation-preserving spin permutations (1,000 rotations). Box plots depict the null distribution of network-mean correlations, where each value reflects the mean correlation across all regions within a given network after spatial rotation. The center line indicates the median, box boundaries represent the interquartile range (25th to 75th percentiles), and whiskers denote the 2.5th and 97.5th percentile. Circles indicate empirical network-mean correlations, with colored filled circles denoting networks whose empirical values exceed the spin-based null distribution and white circles denote non-significant networks. \*\**p* < .01, \**p* < .05. From Base- to Early-learning, greater Limbic eccentricity was associated with better overall learning. From Early- to Late-learning, this pattern reversed: decreased Limbic eccentricity was associated with better overall learning. From Base- to Late-learning, greater DAN contraction was associated with better learning performance. (G–J) Scatterplots depict eccentricity–behavior relationships for representative regions within each significant network (Limbic, SalVentAttn, DMN, DAN). For each network, the region exhibiting the strongest correlation was selected. The top right inset shows seed regions projected onto surface space and colored by network assignment.

We found stage-dependent associations between manifold reconfiguration and cross-task learning performance across multiple higher-order networks. From Base- to Early-learning, we found that increased eccentricity in the Limbic network was selectively associated with better overall learning performance, indicating that a stronger early segregation of limbic regions supported generalized learning ability. No other network-level changes showed meaningful associations with behavioural performance. Notably, from Early- to Late-learning, this brain-behaviour relationship reversed in sign: decreased Limbic eccentricity was now associated with better overall performance, indicating that more successful learners showed dynamic reintegration of limbic systems as learning progressed. During this same phase, decreases in SalVentAttn eccentricity and increases in DMN eccentricity also predicted learning performance. Finally, from Base- to Late-learning, we found that greater contraction within the DAN was associated with better overall learning. Representative regions from each significant network are shown in Fig. 5G–J.

To identify the specific connectivity changes driving these behaviourally relevant manifold effects, we conducted seed-based analyses using representative Limbic and SalVentAttn regions (see Fig. 6A–F; see Supplementary Fig. 6 for DMN and DAN seeds, and Supplementary Figs. 7-8 for complementary seeds in the other hemisphere). Rather than averaging connectivity across each significant network, we selected a single representative seed per network; preserving sensitivity to within-network connectivity changes, such as the limbic segregation-to-integration reversal, that would be obscured by network-level averaging. As above, we present unthresholded maps to illustrate the full spatial distribution of connectivity changes underlying the eccentricity results. We first examined how each seed’s whole-brain connectivity profile changed across learning stages at the group level.

**Figure 6.**
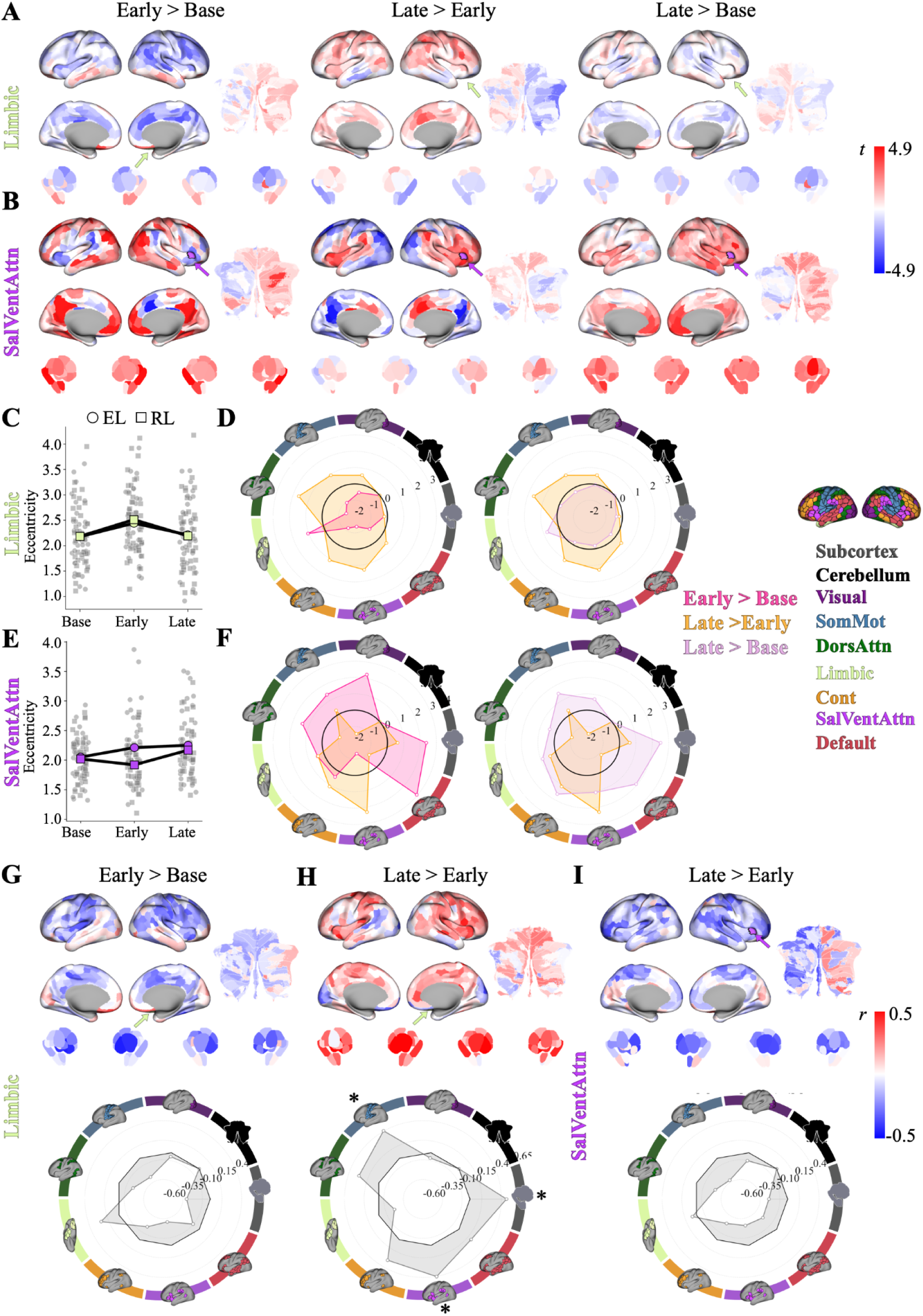
Changes in limbic and attentional connectivity that underlie the relationship between manifold reconfiguration and learning performance. (A–B) Seed-based connectivity changes for representative Limbic (A) and SalVentAttn (B) regions across epoch contrasts (Early>Base, Late>Early, and Late>Base). Red indicates increased connectivity; blue indicates decreased connectivity. Seed regions are indicated by arrows colored by Yeo 7-network assignment (46). For the Limbic seed, Early-learning was characterized by increased within-network connectivity and decreased coupling with distributed systems; this reversed during Late-learning, with increased connectivity to attentional, control, and transmodal networks. For the SalVentAttn seed, Late-learning was characterized by strengthened within-network coupling and increased control network connectivity, alongside disengagement from sensory and DMN systems. (C, E) Mean eccentricity across epochs, for each task (EL, RL) and representative seed region, with individual participants in gray. (D, F) Polar plots summarizing seed-based connectivity changes at the network-level for each contrast. (G–I) Correlations between seed-based connectivity changes and participants’ cross-task learning scores for epoch contrasts reaching network-level significance (from Fig. 5D,E). (G) For the Limbic seed, better learners showed greater Early within-network increases coupled with decreased distributed connectivity. (H) This pattern reversed during Late-learning, with the Limbic seed showing greater connectivity with somatomotor, attentional, DMN, and subcortical networks (I). For the SalVentAttn seed, better learners showed greater late decreases in widespread cortical connectivity, with preserved Limbic coupling. Correlations projected onto surface space (top) and summarized in polar plots (bottom). Maps are presented unthresholded to capture the full spatial distribution of connectivity changes underlying manifold eccentricity effects. Asterisks denote significant correlations following FDR correction \**p* < .05.

For the Limbic seed (see Fig. 6A, C–D), the eccentricity increase from Base- to Early-learning was accompanied by increased coupling with other Limbic regions (i.e. within-network connectivity) alongside decreased connectivity with the DAN, somatomotor, visual, and DMN networks (i.e., limbic segregation). From Early- to Late-learning, this pattern largely reversed: the Limbic seed showed similar within-network connectivity but increased coupling with attention, control, and transmodal networks (i.e., integration), mirroring the stage-dependent changes in the associations with eccentricity change observed in Figure 5a.

For the SalVentAttn seed (see Fig. 6B, E–F), we focused on the Early-to-Late transition, which was the epoch contrast that showed a significant network-level eccentricity–behavior association (see Fig. 5). During this phase, the SalVentAttn seed strengthened its within-network communication and increased coupling with the control and limbic networks. At the same time, connectivity with the DMN, visual, and DAN networks decreased. This pattern is generally consistent with increasing functional segregation and alignment with higher-order control systems as learning plateaued.

### Individual variability in higher-order connectivity changes relate to cross-task learning scores

The analyses above establish that the eccentricity changes in the Limbic and SalVentAttn networks — which we showed are associated with learning performance at the network-level (Fig. 5) — are driven by specific, interpretable redistributions of connectivity across learning stages. What these group-level seed maps cannot tell us, however, is whether the *magnitude* of these connectivity changes varies across individuals in a way that tracks overall learning performance. We addressed this latter question by correlating each participant’s seed-based connectivity changes with their cross-task learning scores for the epoch contrasts that reached network-level significance (see Fig. 6G–I).

For the Limbic seed, the brain-behavior associations closely mirrored the group-level connectivity patterns. Better learners (i.e., higher cross-task learning scores) showed greater early increases in within-network connectivity coupled with greater decreases in connectivity with several distributed networks — most prominently, the SalVentAttn, somatomotor, and subcortical networks (Early>Base; see Fig. 6G). This is the same segregation pattern observed at the group level (noted above), but now linked to individual differences in performance. From Early- to Late-learning, this pattern reversed: better learners now showed greater increases in Limbic connectivity with SalVentAttn, somatomotor, and subcortical networks, alongside decreases in within-network connectivity (see Fig. 6H), largely recapitulating the group-level reintegration pattern. One notable departure was the emergence of stronger Limbic-subcortical coupling in better learners than was apparent in the group-level seed maps (compared Figs. 6G-H vs. Fig. 6A, suggesting that the subcortical component of limbic reintegration may be particularly relevant to individual differences in learning ability.

For the SalVentAttn seed, better learners showed greater decreases in connectivity with widespread cortical networks from Early- to Late-learning, with relative preservation of coupling with the Limbic system (see Fig. 6I). Although this pattern appears to differ from group-level increases in SalVentAttn integration described above, they suggest that the behaviourally relevant component of SalVentAttn reconfiguration may reflect selective decoupling from distributed cortical systems, rather than global integration. Together, these seed-level results indicate that individual differences in learning performance across tasks are associated with flexible, stage-dependent redistributions of connectivity within higher-order limbic and attentional systems — not with modulation of primary sensorimotor circuits.

## DISCUSSION

A dominant framework of motor learning assigns error-based and reinforcement-based processes to separate neural circuits through the cerebellum and basal ganglia, respectively (1, 2, 4, 5). By directly comparing whole-brain activity patterns during both forms of learning within the same participants, we show that this separation is far less pronounced than these models would predict at the whole-brain level. A clear pattern emerged where learning-related network reconfigurations were overwhelmingly shared across EL and RL and concentrated in higher-order transmodal cortex, with task-specific modulation confined to a single exception, in a PMC cluster. Notably, this shared architecture extended to the cerebellum and basal ganglia themselves, the canonical substrates of the EL/RL dichotomy, which reconfigured comparably across both tasks. Even the PMC cluster, the one region to distinguish tasks, did not engage these subcortical structures along classically predicted lines: it coupled with dorsal striatal circuits during EL and with cerebellar circuits during RL. At the level of individual differences, we found that cross-task learning performance was associated not with sensorimotor network changes but with stage-dependent reconfigurations within limbic and higher-order attentional systems. These findings suggest that what distinguishes successful learners is not task-specific circuits but the flexibility with which higher-order networks reconfigure across learning stages and feedback contexts.

The two tasks we used differed substantially in their motor demands. The EL task required discrete reaching movements toward radial targets under rotated visual feedback, whereas the RL task required continuous tracing of a curved path guided solely by scalar reward — differing in movement type, spatial structure, and trial dynamics. That learning-related manifold changes were broadly shared across these motorically distinct tasks suggests that the observed network reconfigurations reflect properties of the learning process itself, rather than common sensorimotor demands. Within this shared architecture, the DMN emerges as a dominant locus: higher-order transmodal cortex rather than task-specific sensorimotor circuits appears to carry the signature of motor learning regardless of feedback type. Several properties of the DMN, documented in other contexts, may be relevant to its role in supporting learning across tasks. The DMN sits near the apex of the cortical processing hierarchy, maximally distant from primary sensory and motor areas (35, 59, 60). This positioning enables broad oversight over distributed brain activity and enables processes — such as strategy formation and performance evaluation — that operate independently of moment-to-moment sensory inputs (61–65). DMN regions also integrate information over extended timescales spanning seconds to minutes (66–68), and allow for the retrieval and reinstatement of previous experiences (69–71), supporting the sustained cognitive effort required for strategy maintenance across trials. Structurally, these regions exhibit lower myelin content, higher neurotransmitter receptor density, and more complex dendritic morphology than sensorimotor cortex (72–74) — features that may confer enhanced capacity for learning, plasticity and computational flexibility (75). Together, these properties suggest that the DMN may provide a cognitive scaffolding that supports both EL and RL.

This interpretation aligns with our recent work showing that both across-day savings and across-hand transfer of visuomotor adaptation are supported by the re-expression of the same DMN manifold structure established during initial learning (40, 76). Our findings extend this principle. The same DMN reconfigurations emerge not just across days and effectors, but across fundamentally different learning processes and tasks (i.e., EL vs. RL). The finding that these motorically and computationally distinct tasks converge on the same transmodal network architecture supports a shared role for the DMN across different forms of motor learning. This interpretation is further reinforced by the observation that the majority of learning-related manifold changes did not simply revert to baseline as learning performance plateaued. Although a small subset of regions returned to their pre-learning connectivity state, most retained altered eccentricity profiles through Late-learning — and this persistence was shared across both tasks, despite the fact that behavioral errors in EL had returned to near-baseline levels and that directional error signals were never present in RL. The combination of largely absent error signals and cross-task convergence makes ongoing error processing an unlikely driver of these late-learning connectivity patterns, and implicates a persistent neural state associated with the acquired motor adaptation.

This persistent, task-general signature is particularly evident in the subcortical and cerebellar structures at the heart of the classical EL/RL dichotomy. As noted, Dual-systems models hold that these structures should dissociate across tasks — the cerebellum preferentially engaging during EL, the basal ganglia during RL (1, 4, 5, 11). Our data do not support this at the level of whole-brain network organization. Instead, cerebellar and striatal regions reconfigured comparably across both tasks, along shared trajectories, and with no regions in either structure showing a significant Task or Task × Epoch interaction effect. The cerebellar reconfigurations we observed were themselves non-monotonic, with widespread early expansion giving way to selective contraction of cognitive/association subdivisions as learning progressed — a dynamic profile inconsistent with a narrow view of the cerebellum as primarily an error-correction structure (5–7, 11, 22–24). These findings align with recent evidence that cerebellar circuits participate in reward-based learning (25–28) and that the functional contributions of cortico-cerebellar and cortico-striatal circuits to learning are less cleanly dissociated than classical models suggest (27, 45). Against this shared subcortical backdrop, we nevertheless find that higher-order cortex selectively couples to functional subcircuits within the cerebellum and basal ganglia depending on the learning context — a pattern we examine next through the PMC’s task-specific coupling.

At the task level, the PMC selectively altered its connectivity depending on the type of learning signal provided, coupling with striatal and action-related cerebellar circuits during EL but with hippocampal and cognitive/association cerebellar circuits during RL. These couplings invert the subcortical partners that classical models would place with each learning type: the caudate and putamen — typically associated with RL (1, 4) — emerged as strong PMC partners during EL, while the cerebellum — typically associated with EL (5, 6, 11, 22–24) — emerged as a strong PMC partner during RL. Within the cerebellum, this RL-aligned coupling was further specific to association/cognitive subregions rather than action-related motor zones, providing a finer level of circuit specificity within a structure traditionally associated with EL. At the epoch level, the PMC disengaged from its parent DMN at the onset of learning to interface with task-specific sensorimotor circuits, then gradually reintegrated with broader association cortex as performance stabilized. Notably, the PMC was the only set of regions in the brain to exhibit both task and epoch sensitivity — a dual flexibility across feedback type and learning stage that positions it, and particularly the PCC subregions within it, as a site of convergence for learning-related network dynamics. Whether the PMC actively coordinates these reconfigurations or instead reflects their downstream integration cannot be resolved from the present data, and will require directed connectivity approaches to address. However, this convergent role aligns with the PCC’s established characterization as a cortical hub that contains “echoes” of multiple brain networks within its subregions (77, 78), and with its dense structural connectivity to virtually every major brain system (77). These properties may be what underlies the PMC’s capacity to differentially express connectivity patterns across both distinct learning contexts and stages.

Another notable finding concerns the neural basis of individual differences in learning performance across tasks. We found that participants who tended to perform well across both tasks showed distinctive patterns of limbic network reconfiguration — a result that, at first glance, seems unexpected given that limbic regions have not commonly been associated with *motor* learning. Yet the limbic cortex shares, and in several respects exceeds, the very properties that position the DMN for a role in learning. Like the DMN, limbic regions sit at the apex of cortical processing hierarchies (29, 35, 79), but they represent the most extreme end of this gradient: they exhibit the simplest laminar architecture in the cerebral cortex (agranular and dysgranular) and the lowest intracortical myelin content of any cortical zone (30, 72, 80, 81). Whereas the DMN’s contribution appears to operate at the group level (shared transmodal reconfigurations accompanying learning regardless of feedback type), the limbic network’s contribution is individual-specific and stage-dependent. During Early-learning, better learners exhibited greater limbic segregation, involving increased within-network connectivity and decreased coupling with distributed systems. However, as learning progressed, this pattern reversed — better learners showed greater limbic reintegration with somatomotor, attentional, DMN, and subcortical networks. This reversal is inconsistent with a monotonic relationship between network integration and learning performance. Instead, it indicates that successful learners flexibly modulate the limbic network’s coupling with distributed systems as a function of the learning stage. Early segregation may serve to consolidate or protect nascent learning representations (82), while later reintegration enables broader network coordination as motor performance stabilizes. This dynamic profile is consistent with a system that initially shields fragile learning signals from interference (83, 84) and later broadcasts them across distributed networks (85) to guide coordinated motor refinement. The limbic finding was not isolated — the SalVentAttn, DMN, and DAN networks also showed significant stage-dependent associations with learning performance, reinforcing the conclusion that learning performance across tasks is associated with the flexible reconfiguration of higher-order networks rather than modulation of primary sensorimotor circuits.

### Methodological Considerations

One concern common to all learning studies is that any observed neural changes may reflect nonspecific temporal factors (e.g., habituation, fatigue, or even scanner drift) rather than genuine learning-related dynamics. Several features of our data argue against this interpretation. The manifold changes we observed were non-monotonic across both cortical and subcortical structures; i.e., eccentricity in many regions increased from Base to Early-learning but reversed from Early to Late-learning. This was evident in the cerebellum and amygdala as well as in the transmodal cortex — a pattern inconsistent with any monotonic time-dependent process. The limbic network’s relationship with behavior provides particularly direct evidence: better learners tended to show increased segregation early and increased integration late, a non-monotonic pattern that is difficult to explain by monotonic changes in attention or arousal.

A second consideration is that the RL task was always performed before the EL task. This was a deliberate design choice, as the visuomotor rotation task used in EL can induce exploratory behaviour and explicit re-aiming strategies (16–18, 86) that could alter how participants approach the (subsequent) RL task. Because session order was not counterbalanced, differences between EL and RL — in particular, the task-dependent PMC connectivity patterns — may not be uniquely attributed to feedback type, and some contribution from order-related factors (e.g., scanner familiarity, cumulative practice with the touchpad) cannot be ruled out. Several features of our data nonetheless constrain this interpretation. First, our primary findings concern epoch effects that are expressed within each session, against that session’s own baseline, and are therefore unaffected by between-session confounds. Second, the task effects that did emerge were anatomically focal rather than global: only 11 regions, concentrated within a single cluster in the PMC, showed significant task effects, against 301 regions showing epoch effects. A nonspecific shift in brain state between sessions would be expected to produce far more distributed task-related differences. Still, replication with a counterbalanced design will be required to confirm that the PMC’s differential coupling with striatal versus cerebellar-hippocampal circuits is specific to sensory feedback type per se.

A final consideration concerns the behavioral index that we used to relate manifold reconfigurations to individual differences in learning. Our cross-task learning score (fPC1) was significantly associated with performance in both EL and RL, but captured a greater share of behavioral variance in RL (r = 0.99) than in EL (r = 0.33). This asymmetry likely reflects the greater inter-individual variability found in the RL task (see Fig. 2), where the absence of directional error information places greater demands on strategic search and produces wider variation in learning trajectories (18, 21, 39). That EL-only fPC1 scores nonetheless correlated significantly with the combined shared dimension is consistent with the cross-task learning score capturing a behavioral axis that spans both tasks rather than one unique to RL — indicating that participants who performed well under reinforcement feedback also tended to perform well under error-based feedback. This is despite the two tasks differing in feedback type, motor structure, and being separated by a one-week interval between scanning sessions. As the shared learning dimension is expressed more variably in RL, the brain-behavior patterns we report are best understood as reflecting a dimension that captures performance in both tasks, but that RL resolves with greater sensitivity.

### Conclusion

Classical models of motor learning assign error-based and reinforcement-based processes to anatomically separate neural circuits. The current findings demonstrate that these forms of learning converge on a shared functional architecture spanning transmodal cortex, limbic, and subcortical structures — and what distinguishes successful learners is not changes in task-specific circuits but the flexibility with which higher-order networks reconfigure across learning stages and feedback contexts. By situating motor learning within the broader framework of large-scale network dynamics, our work highlights a prominent role for transmodal cortical systems in supporting flexible adaptation. Understanding motor learning, and the individual differences that characterize it, may depend less on the regions that compute error or reward signals and more on the distributed, higher-order network dynamics that coordinate them.

## MATERIALS & METHODS

Forty-six right-handed individuals (27 female, aged 18–28 years) completed three sessions: a mock-scanner training session, followed by an RL and an EL MRI session, each approximately one week apart. Nine participants were excluded for excessive head motion, scan interruption, incomplete trials, or failure to follow task instructions, yielding a final sample of 37 individuals. The Queen’s University Research Ethics Board approved the study and it was conducted in accordance with the principles outlined in the Canadian Tri-Council Policy Statement on Ethical Conduct for Research Involving Humans and the Declaration of Helsinki (1964).

In the RL task, participants performed a hidden-reward tracing task in which they attempted to trace a curved path without online visual feedback of their finger trajectory (see Fig. 1D). Although participants were told performance scores reflected how accurately they traced the visible path, reward score was actually based on a horizontally mirrored version of the visible path. Thus, learning depended on reward-based trial and error rather than sensory prediction error (20). In the EL task, participants completed a classic visuomotor rotation paradigm (7, 16, 17) where they made center-out movements to visual targets. After baseline trials with veridical cursor feedback, a 45° clockwise rotation was applied to the cursor, allowing us to measure adaptation to sensory prediction errors.

Behavioral performance was quantified separately for each task and combined into a single measure of overall learning performance using functional principal components analysis (fPCA; <u>42</u>). Learning trajectories from both tasks were concatenated and submitted to fPCA, yielding a set of functional components that captured individual differences in performance over time. The first component (fPC1) explained 61.2% of the variance and served as our primary behavioral index — hereafter referred to as the cross-task learning score — with higher scores indicating better learning across tasks.

MRI data were acquired using a 3-Tesla Siemens TIM MAGNETOM Trio MRI scanner located at the Centre for Neuroscience Studies, Queen’s University. Imaging data were preprocessed using fMRIprep 20.1.1 (87). We extracted regional BOLD timeseries data for cortical parcels derived from the Schaefer 400-parcellation (46), subcortical regions predefined according to the Tian scale II, 3T subcortical atlas (47), and cerebellar regions defined by the Nettekoven 32-region cerebellar atlas (48). For each participant, task-based scans were divided into six equal-length epochs: Base, Early, and Late-learning for both the RL and EL tasks (198 imaging volumes each). For each participant, task, and epoch, we estimated FC matrices using Ledoit-Wolf covariance estimation, and applied a Riemannian centering approach to remove subject-level variance in covariance structure (37–39, 51, 88).

We next used manifold learning to characterize large-scale changes in network organization across learning. For each participant, task, and epoch, covariance matrices were row-wise thresholded and converted into cosine similarity affinity matrices. We applied PCA to each of these matrices to generate low-dimensional representations of whole-brain FC. We then quantified each region’s position within this low-dimensional manifold space using manifold eccentricity, defined as the Euclidean distance from the manifold centroid. Higher eccentricity reflects a more segregated connectivity profile, whereas lower eccentricity reflects greater integration with other brain networks (35–40).

Regional eccentricity values were analyzed using a rmANOVA with Task (EL, RL) and Epoch (Base, Early, Late) as within-subject factors, followed by paired-samples t-tests across significant regions for each comparison. To determine whether these manifold changes were related to behavior, we correlated participants’ cross-task learning scores with eccentricity changes across key epoch contrasts (Early>Base, Late>Early, and Late>Base). Network-level effects were assessed using the Yeo 7-network parcellation and spin permutation testing. To characterize the specific connectivity changes underlying significant manifold effects, we performed seed-based connectivity analyses using representative regions from each significant network, summarized at both the whole-brain voxel level and aggregated at the network-level. Full details are provided in *SI Materials and Methods*.

