## Supplemental Information for "A shared functional architecture for error-based and reinforcement-based motor learning in the human brain"

**Supplemental Figure 1:** fPC1 scores are robust to RL trial truncation

**Supplemental Figure 2:** Combined fPC1 scores capture individual differences in learning ability within each task separately

**Supplemental Figure 3:** A clear inflection point at five components supports dimensionality selection for the connectivity manifold

**Supplemental Figure 4:** Manifold eccentricity tracks canonical graph-theoretic measures of functional segregation and integration

**Supplemental Figure 5:** Distribution of F-values across effects in the rmANOVA

**Supplemental Figure 6:** DMN and DAN seed regions show stage-dependent connectivity changes that parallel the limbic and attentional patternsd

**Supplemental Figure 7:** Contralateral limbic and SalVentAttn seed regions replicate the stage-dependent connectivity patterns observed in the primary seeds

**Supplemental Figure 8:** Contralateral DMN and DAN seed regions replicate the stage-dependent connectivity patterns observed in the primary seeds

**Supplemental Figure 9:** Task and epoch effects on manifold eccentricity are stable across a range of dimensionality choices.

**Supplemental Figure 10:** Learning-related manifold changes are not driven by differences in mean BOLD signal amplitude

#### Detailed Materials and Methods

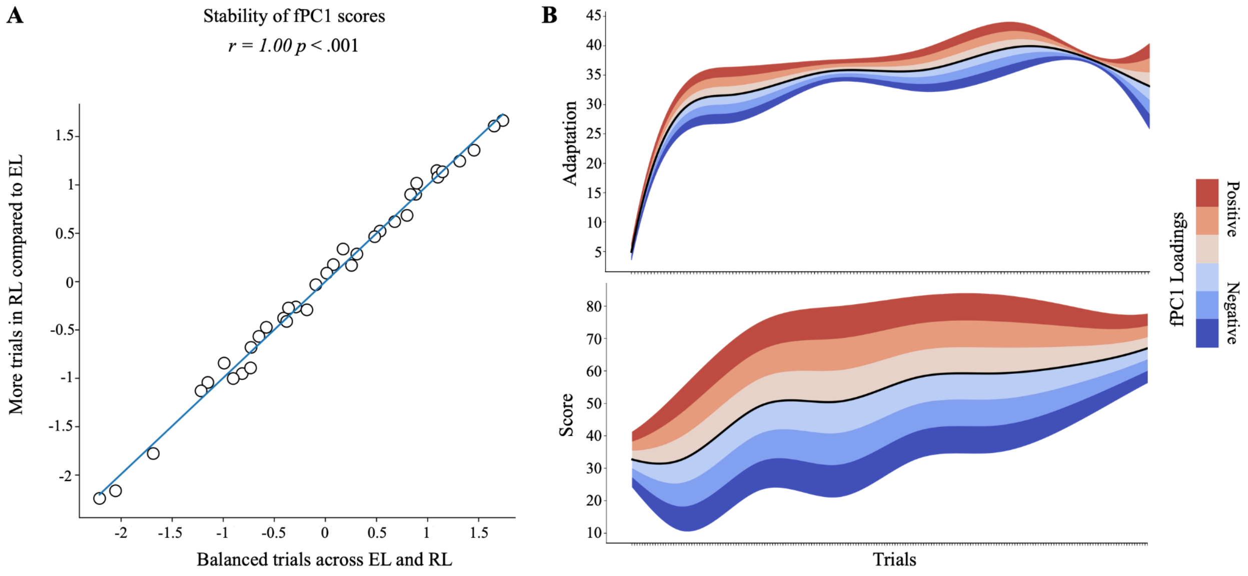


**Supplemental Figure 1. fPC1 scores are robust to RL trial truncation.** (A) Scatter plot demonstrating the correspondence between participants' fPC1 scores computed using a truncated (160 trials) versus the full (200 trials) RL learning trajectories. (B) Learning curves for the EL task, and the RL task across trials, showing the full time course of performance used in our validation analysis.

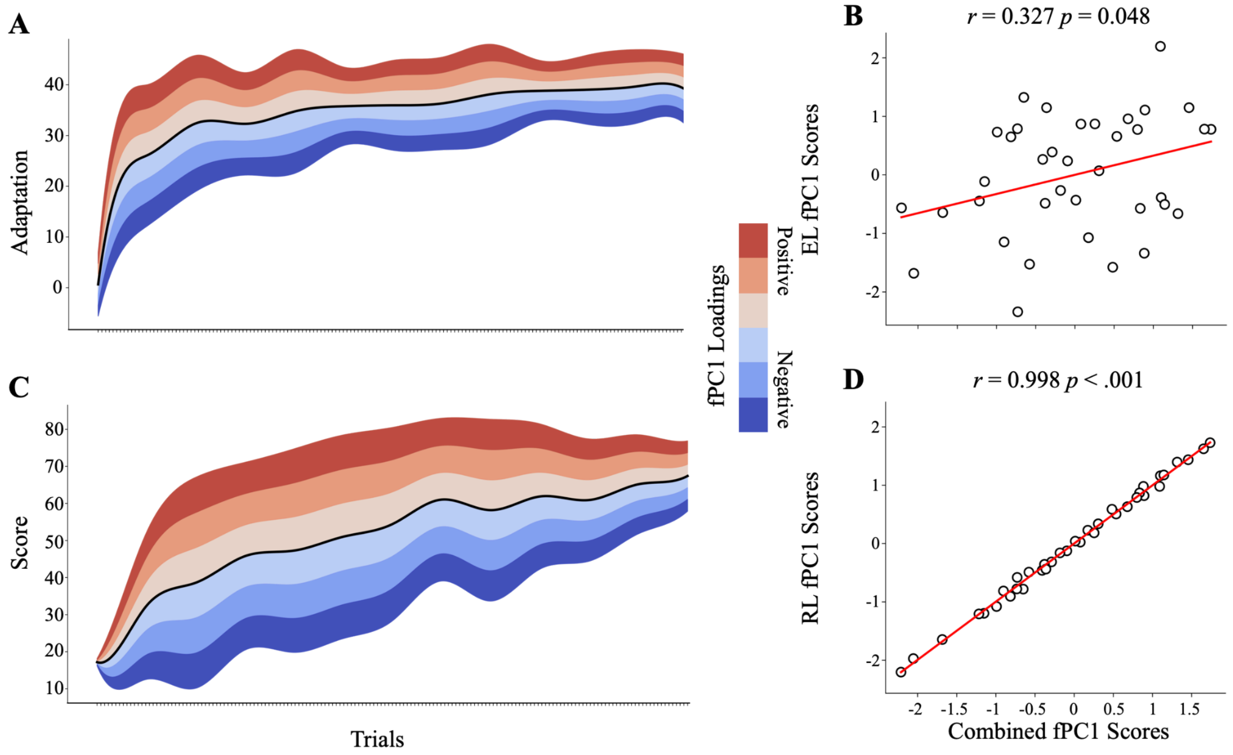


**Supplemental Figure 2. Combined fPC1 scores capture individual differences in learning ability within each task separately.** (A) fPCA applied to the EL adaptation curves along, showing the effect of positive (red) and negative (blue) task-specific fPC1 scores relative to the mean (black). (B) Correlation between the combined (cross-task) fPC1 scores (used in the main paper; see Main Paper Fig. 1G) and the EL-only fPC1 score, demonstrating that the shared dimension captures individual differences in error-based adaptation. (C) fPCA applied to RL score curves alone. (D) Correlation between the combined fPC1 score (used in the main paper; see Main Paper Fig. 1G) and the RL-only fPC1 score, demonstrating that the shared learning dimension also captures individual differences in reinforcement-based learning. Significant positive correlations in both (B) and (D) confirm that the combined fPC1 is not purely dominated by either task alone, but captures a common axis of learning variability expressed across both EL and RL.

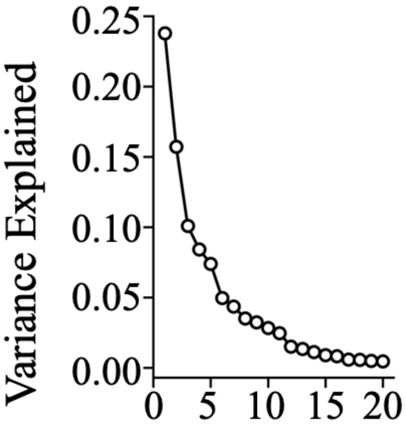


**Supplemental Figure 3. A clear inflection point at five components supports dimensionality selection for the connectivity manifold.** The percent variance explained by each of the first 20 principal components from their functional connectivity PCA. Together, the top five PCs account for 65.4% of total variance.

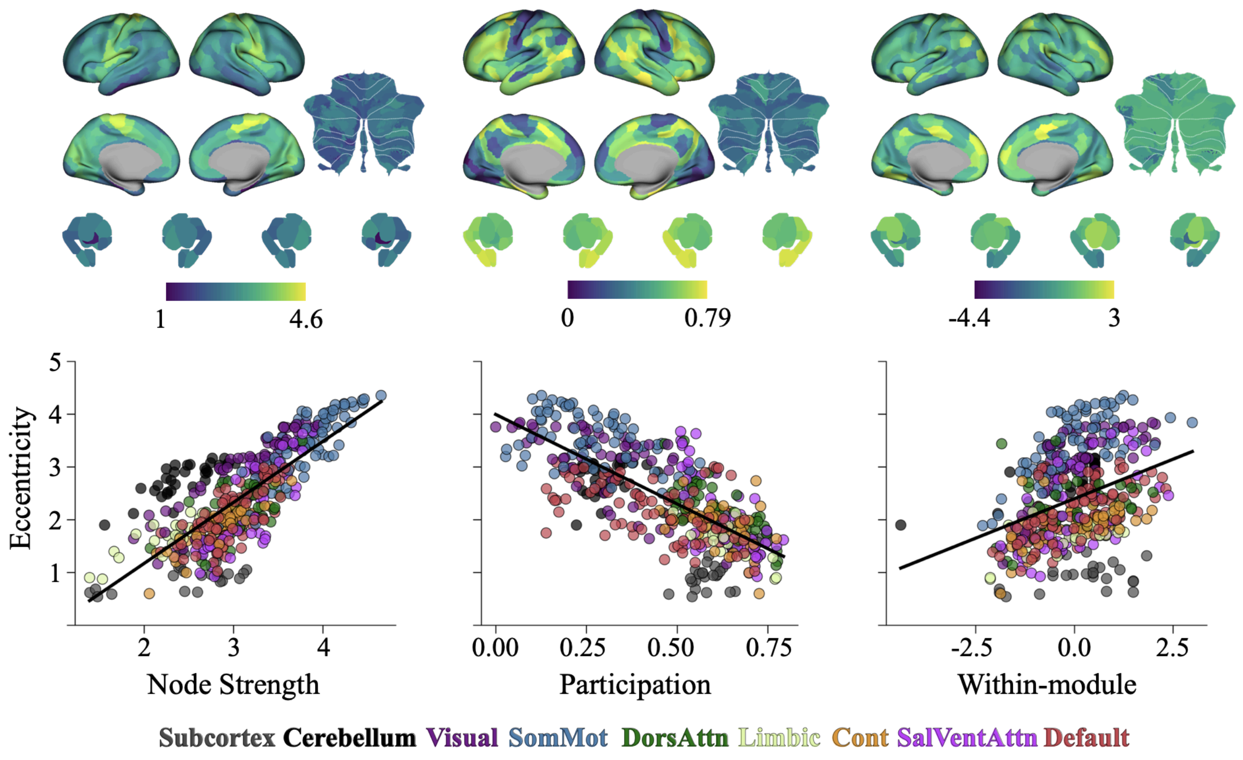


**Supplementary Figure 4. Manifold eccentricity tracks canonical graph-theoretic measures of functional segregation and integration.** *Top,* maps of node strength, participation coefficient, and within-module degree z-score for the group-average resting connectivity matrix (i.e., reference connectivity matrix). *Bottom,* corresponding correlations between each functional connectivity measure and manifold eccentricity, coloured according to the Yeo 7-network parcellation (32), with the subcortex in grey (33) and cerebellum in black (34).

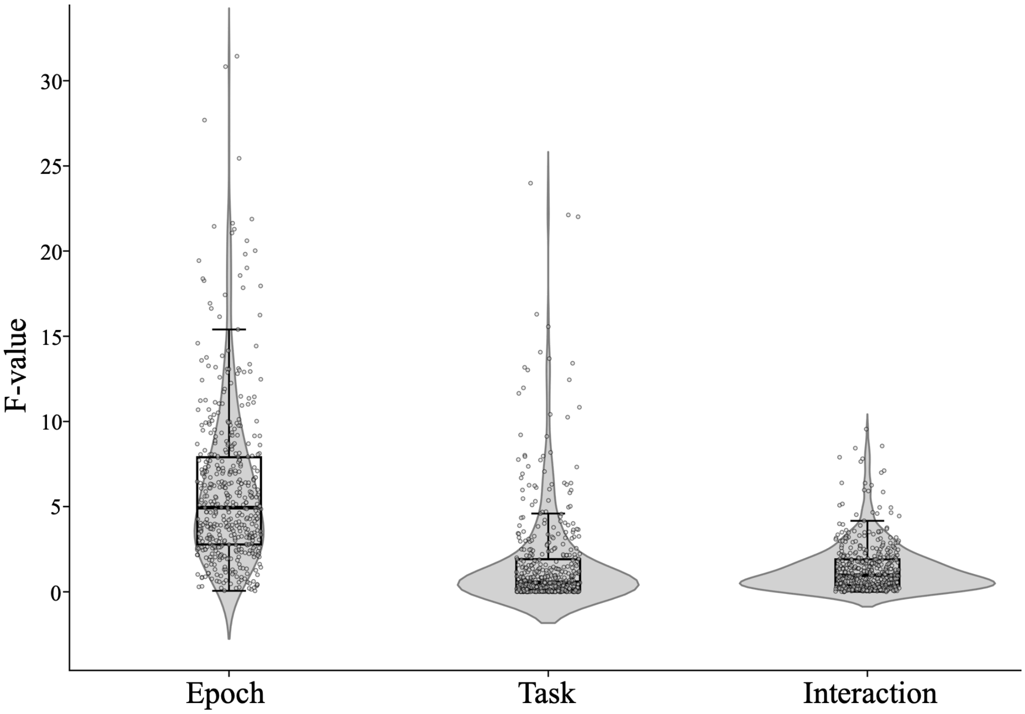


**Supplementary Figure 5. Distribution of F-values across effects in the rmANOVA.** Violin plots show the distribution of F-values for the main effects of Epoch (Base, Early, Late) and Task (EL, RL), as well as their interaction (Task × Epoch), across all brain regions. Epoch effects were substantially larger than both Task and interaction effects, confirming that the design has clear sensitivity to detect learning-related network reconfigurations. Notably, the distribution of interaction F-values closely resembled that of the Task main effect, indicating that interaction effects were not merely underpowered versions of large true effects, but were comparable in magnitude to the minimal task-related modulation observed. Box plots indicate median and interquartile range; scatter points represent individual regions.

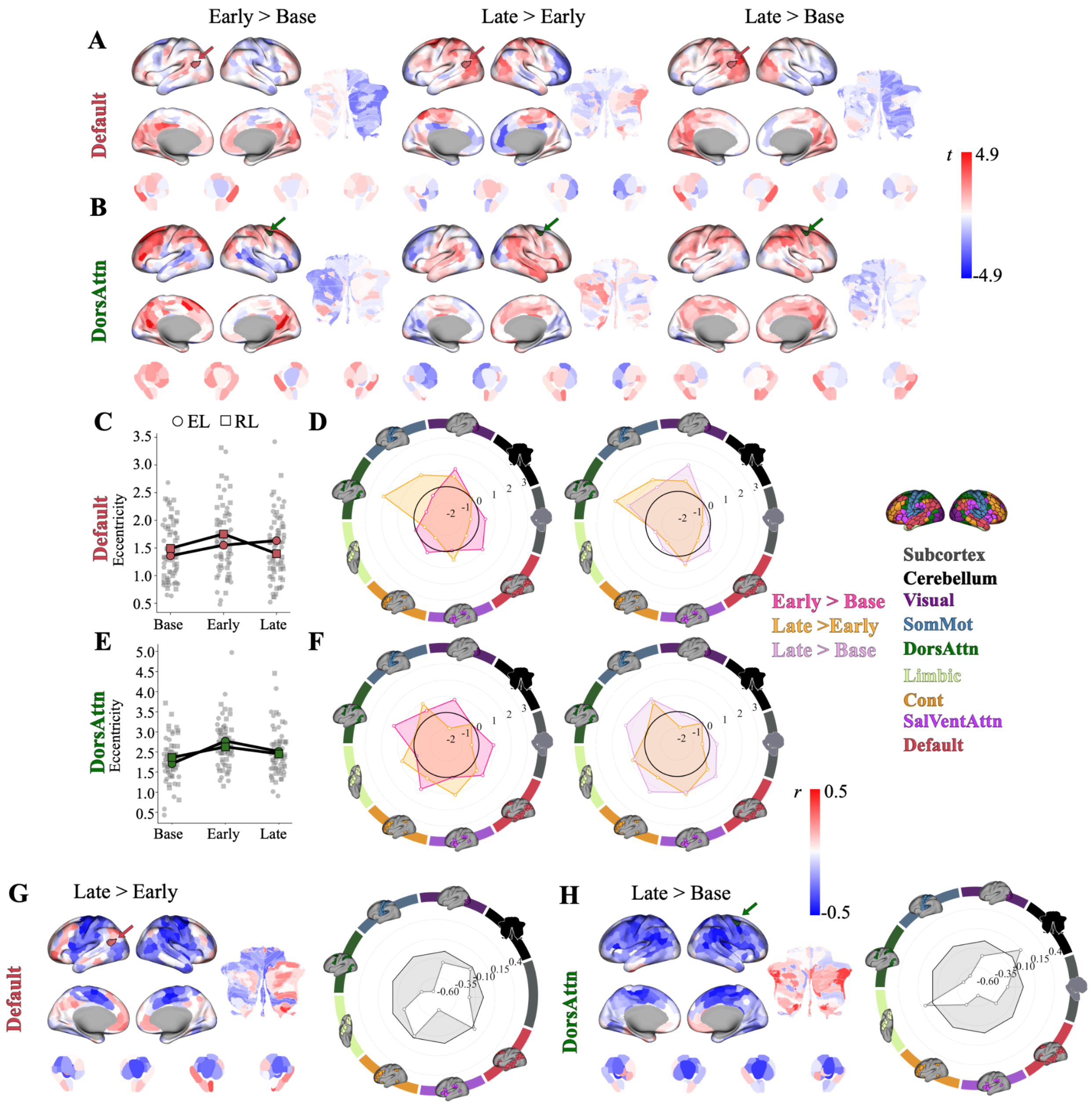


**Supplemental Figure 6. DMN and DAN seed regions show stage-dependent connectivity changes that parallel the limbic and attentional patterns.**(A-B) Connectivity changes of representative brain regions in the Default (A) and Dorsal Attention (B) network from Early > Base, Late > Early, and Late > Base learning. Positive (red) values show increases and negative (blue) values show decreases in connectivity. Each seed region is indicated by their functional colour assignment according to the Yeo 7-network parcellation, as indicated by the arrows in their respective colours. (C, E) Scatterplots show representative seed regions mean eccentricity for each participant across epochs, with individual participant points in light grey. (D, F) Polar plots show seed-based connectivity changes between epochs at the network level (according to the Yeo 7-network parcellation, as well as the addition of the subcortex and cerebellum). The colour behind each brain region indicates its functional network assignment, with the polar plots indicating the colour of the epoch contrast. (G-H) Seed connectivity difference correlations against epochs of interest for Default and Dorsal Attention network representative regions, respectively, and fPC1 scores. Correlations for each network, aggregated based on their functional network assignment (according to the Yeo 7-network parcellation (32), as well as the addition of the subcortex (33) and cerebellum (34). Correlations are projected in both surface space on the left, and in polar plots on the right. Each seed region is indicated by their functional colour assignment according to the Yeo 7-network parcellation, as indicated by the arrows in their respective colours. The colour behind each brain region indicates its functional network assignment, with the polar plots indicating the colour of the task or epoch contrast (see right).

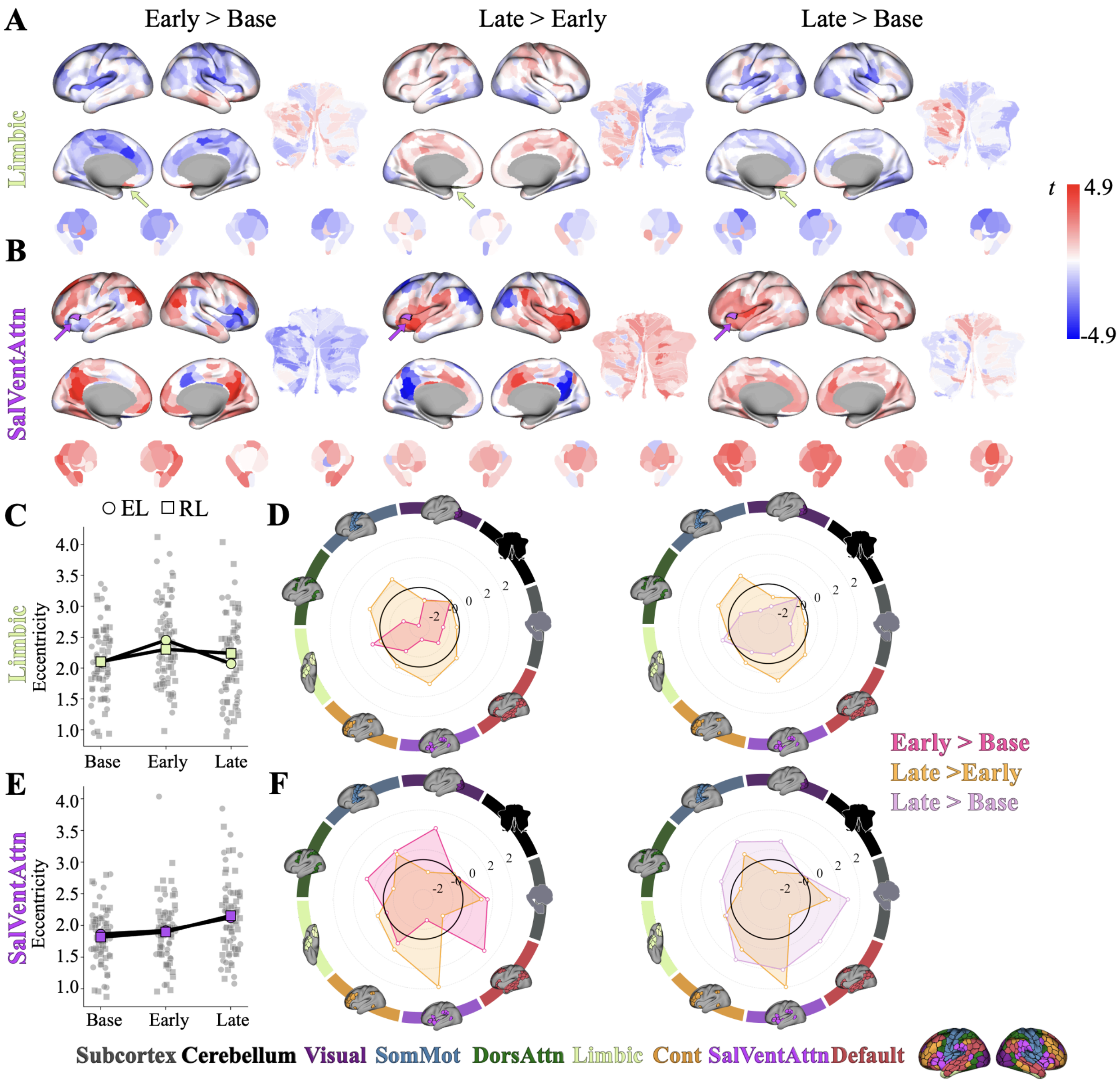


**Supplemental Figure 7. Contralateral limbic and SalVentAttn seed regions replicate the stage-dependent connectivity patterns observed in the primary seeds.** (A-B) Connectivity changes of representative brain regions in the Limbic (A) and SalVent Attention (B) network from Early > Base, Late > Early, and Late > Base learning. Positive (red) values show increases and negative (blue) values show decreases in connectivity. Each seed region is indicated by their functional colour assignment according to the Yeo 7-network parcellation, as indicated by the arrows in their respective colours. (C) Scatterplots show representative seed regions mean eccentricity for each participant across epochs, with individual participant points in light grey. (D) Polar plots show seed-based connectivity changes between epochs at the network level (according to the Yeo 7-network parcellation, as well as the addition of the subcortex and cerebellum). The colour behind each brain region indicates its functional network assignment, with the polar plots indicating the colour of the epoch contrast.

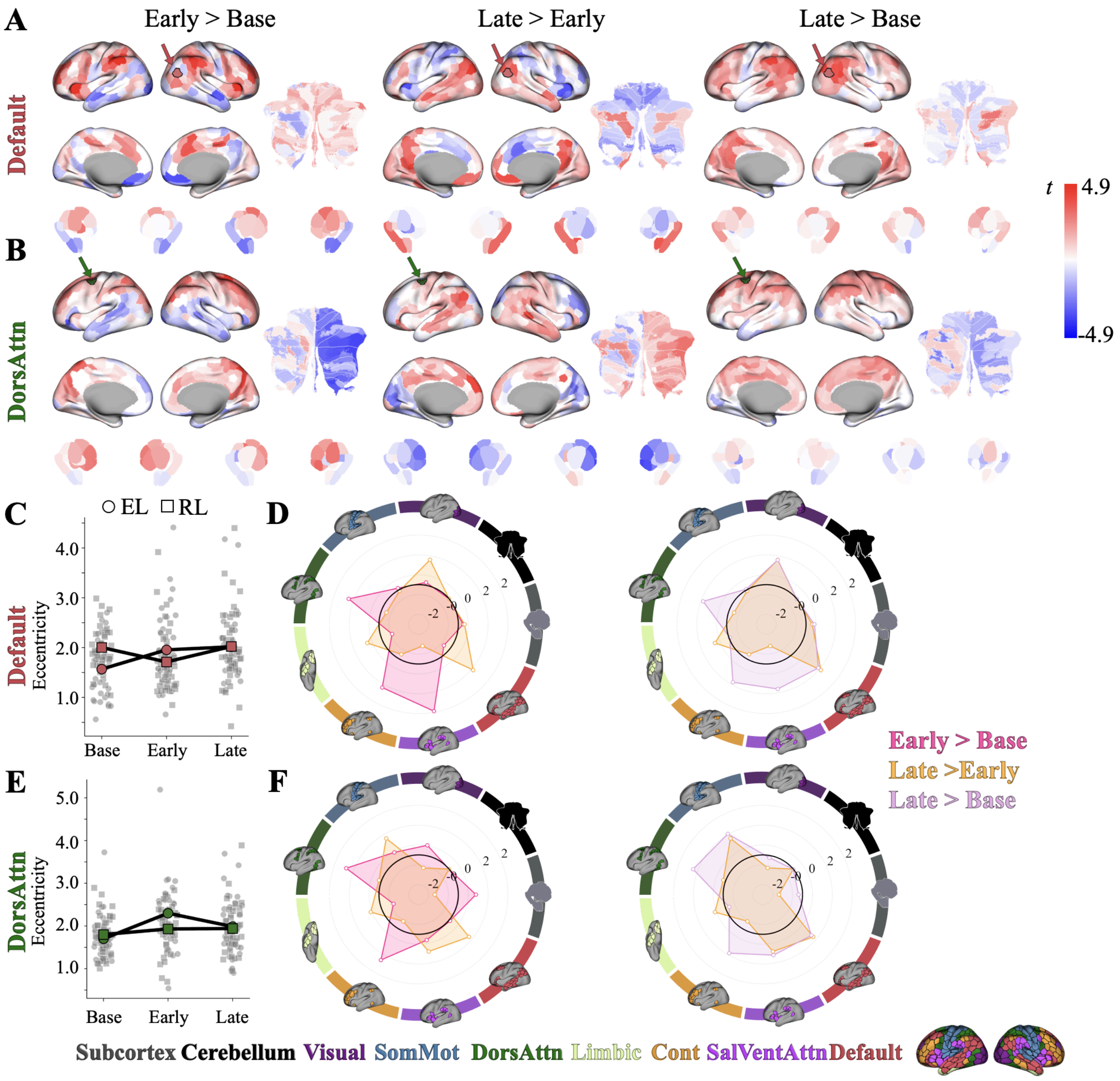


**Supplemental Figure 8. Contralateral DMN and DAN seed regions replicate the stage-dependent connectivity patterns observed in the primary seeds.** (A-B) Connectivity changes of representative brain regions in the Default (A) and Dorsal Attention (B) network from Early > Base, Late > Early, and Late > Base learning. Positive (red) values show increases and negative (blue) values show decreases in connectivity. Each seed region is indicated by their functional colour assignment according to the Yeo 7-network parcellation, as indicated by the arrows in their respective colours. (C) Scatterplots show representative seed regions mean eccentricity for each participant across epochs, with individual participant points in light grey. (D) Polar plots show seed-based connectivity changes between epochs at the network level (according to the Yeo 7-network parcellation, as well as the addition of the subcortex and cerebellum). The colour behind each brain region indicates its functional network assignment, with the polar plots indicating the colour of the epoch contrast.

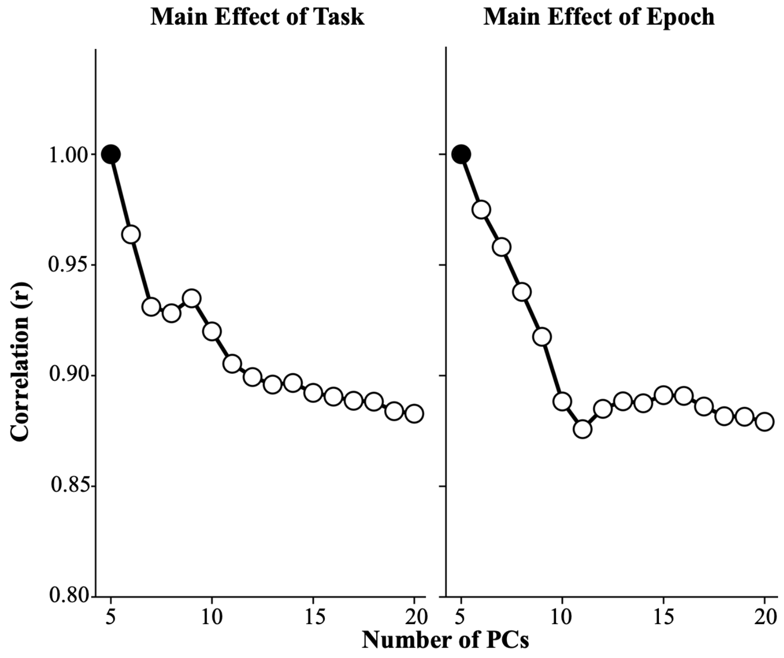


**Supplemental Figure 9. Task and epoch effects on manifold eccentricity are stable across a range of dimensionality choices.** (A-B) Scatter plots show the correlation of F-values for the main effect of (A) Task and (B) Epoch when manifold eccentricity was computed using principal components 5-20. Each point reflects the correlation between F-values obtained with a given number of components and those from our main analyses using five components. The black dot indicates an r-value of 1 (i.e., perfect correspondence with our main analyses), which serves as the reference for comparison. Note that all correlation values are $r > 0.85$, which indicates high reliability.

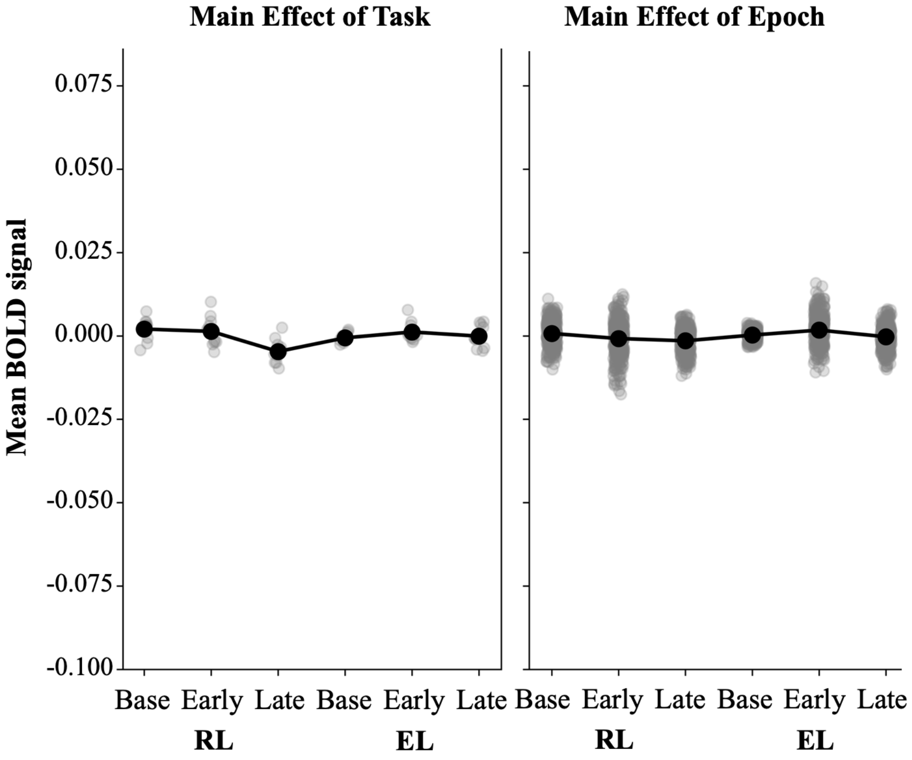


**Supplemental Figure 10. Learning-related manifold changes are not driven by differences in mean BOLD signal amplitude.** Scatter plots depict mean BOLD signal for Task (EL and RL) and Epoch (Base, Early, Late), averaged across ROIs exhibiting significant maine effects in our eccentricity analysis. Grey points represent the mean BOLD signal for individual ROIs, while black dots indicate the average across ROIs for each condition. To test whether differences in mean BOLD signal amplitude could account for the observed eccentricity effects, we performed a Task (EL, RL) x Epoch (Base, Early, Late) repeated measures ANOVA for each ROI, followed by FDR-correction across ROIs for each effect. A main effect of epoch was observed in a subset of the ROIs (54 regions; 17.9% of the original 301 ROIs), while no widespread main effect of Task was detected (2 regions; 18.2% of the original 11 ROIs) and no significant interactions were found. Thus, while our eccentricity analysis revealed widespread main effects of epoch (301 regions) and a subset of ROIs exhibiting significant task effects (11 regions), mean BOLD signals showed far fewer epoch-related effects and minimal task-related effects. This dissociation suggests that the observed connectivity differences are not driven by changes in mean BOLD signal amplitude.

### SUPPORTING TEXT

#### MATERIALS & METHODS

All imaging data were preprocessed using fMRIprep, which is open source and freely available. Our analysis and links to the data are available at: https://github.com/thelamplab/biManifold. All behavioral and imaging data (including T1w and functional scans) have been made freely available at the following repository at OpenNeuro (https://openneuro.org/datasets/ds005598).

##### *Participants*

Forty-six right-handed individuals (27 females, aged 18–28 years, M = 20.3) were recruited and participated in three separate testing sessions: the first, an MRI-training session, was followed approximately one week later by the RL MRI session, and approximately one week after that by the EL MRI session. The RL session was always conducted before the EL session, to avoid potential contamination from explicit re-aiming strategies induced by the visuomotor rotation task. Of these 46 participants, 9 were excluded from the final analysis: 1 due to excessive head motion in the scanner (motion greater than 2 mm or 2° rotation within a single scan), 1 due to interruption of the scan during the learning phase of the RL task, 4 due to incomplete trials (>25% of trials not completed within the maximum trial duration in one or both tasks), and 3 due to a failure to properly perform the RL task (these participants did not trace the visible rightward path during the baseline phase, but rather continued to trace a straight line for the entire duration of the task, suggesting that they did not attend to the task instructions). This yielded a final sample of 37 participants. All participants were right-handed according to the Edinburgh handedness questionnaire (1). Participant consent was obtained before beginning the experimental protocol. The Queen's University Research Ethics Board approved the study and it was conducted in accordance with the principles outlined in the Canadian Tri-Council Policy Statement on Ethical Conduct for Research Involving Humans and the principles of the Declaration of Helsinki (1964).

##### *Procedure*

The first session consisted of an MRI-training session inside a mock (0T) scanner, which was made to look and sound like a real MRI scanner. This training session served to familiarize participants with the experimental setup and the two motor learning tasks (EL and RL) that would later be performed during fMRI scanning, as well as to ensure that participants were able to remain still in the scanner environment without discomfort or claustrophobia. To establish baseline performance for each task, participants completed approximately 80 trials per task during the training session. To encourage minimal head movement, participants' head position was monitored in real time using a Polhemus motion tracking sensor (Polhemus, Colchester, VT) affixed to the forehead with medical tape. This system provided continuous measurements of head translation and rotation across six degrees of freedom. When head displacement exceeded 0.5 mm of translation or 0.5° of rotation (within a predefined velocity criterion), participants received an auditory tone delivered through a speaker positioned near the head. This feedback enabled participants to learn to minimize head movement during task performance. Following completion of the training session, participants returned approximately one week later for the RL MRI session, and approximately one week after that for the EL MRI session (see below for details).

##### *Apparatus*

During testing in the mock scanner, participants performed hand movements by applying fingertip pressure to a digitizing touchscreen tablet (Wacom Intuos Pro M). During the MRI sessions, participants used an MRI-compatible digitizing tablet (Hybridmojo LLC, CA, USA). In both the mock and MRI scanners, the target and cursor stimuli were rear-projected onto a screen mounted behind the participant using an LCD projector (NEC LT265 DLP projector: 1024 x 768 resolution; 60 Hz refresh rate). Visual stimuli were viewed via a mirror affixed to the MRI head coil positioned above the participant's eyes, which prevented direct vision of the hand during task performance. Throughout both testing sessions, eye movements were monitored by the experimenter to ensure participants remained attentive to the task.

##### *Reinforcement-based learning (RL) task*

In the RL task, participants learned through reward-based trial-and-error to produce finger movement trajectories matching an unseen target shape. On each trial, participants were instructed to trace a subtly curved path displayed on the screen without visual feedback of their actual finger trajectory (see Main Paper Fig. 1D). Participants were informed that they would receive a score (0–100 points; linked to financial incentive) reflecting how accurately they traced the visible path and were instructed to maximize their score across trials. However, unbeknownst to participants, the reward score was based on a horizontally mirrored version of the path. As they received no visual feedback on their cursor position, learning relied on reinforcement signals rather than sensory prediction errors (for a similar task, see 2).

Each trial began with participants moving a cursor (3 mm radius cyan circle) into a start position (4 mm radius white circle) located at the bottom of the screen. This cursor was visible only within 30 mm of the start position. After holding within the start position for 0.5 s, the cursor disappeared and a curved path and target (horizontal red line; 30 x 1 mm) appeared 60 mm above the start location. The path was a half-sine wave with an amplitude equal to 0.15 x the target distance. Participants were instructed to trace the displayed path. When they reached the target distance, the target changed from red to green to indicate trial completion. Trials terminated after 4.5 s, followed by a 1.5 s inter-trial interval.

During the learning block, performance was scored from 0–100 points. The cursor's x-position was interpolated at fixed y-displacements (10–60 mm increments), and absolute deviations from the rewarded (mirrored) path were summed. This error was normalized relative to a reference sine wave (amplitude = 0.5 x target distance) and scaled to produce a score between 0 and 100. During Base trials (70 trials), no performance feedback was provided. During learning (201 trials), participants received trial-wise scores and were instructed to maximize them. During mock MRI training sessions (i.e., one week prior to the MRI testing session), participants only performed a practice block in which they traced a straight line with (40 trials) and then without (40 trials) visual feedback of the cursor position during the reach. This training session exposed participants to key features of the task (e.g., use of the touchscreen tablet, trial timing) and allowed us to establish adequate performance levels while not allowing for any reward-based learning to take place.

At the beginning of the MRI testing session, but prior to the first scan being collected, participants re-acquainted themselves with the RL task by first performing a practice block in which they traced a straight line with (40 trials) and then without (40 trials) visual feedback of the cursor position during the reach. Next, we collected an anatomical scan, followed by a DTI scan, followed by a resting-state fMRI scan. During the resting-state scan, participants were instructed to rest with their eyes open while fixating on a central cross presented on the screen. Following this, participants performed the RL task, which consisted of two separate experimental runs without visual feedback of the cursor: (1) a baseline block of 70 trials in which they attempted to trace the curved path and no score was provided, and (2) a separate learning block of 200 trials in which participants were instructed to maximize their score shown at the end of each trial.

##### *Error-based learning (EL) task*

To probe EL, we used a well-characterized visuomotor rotation (VMR) paradigm (3–5). During the task, participants performed blocks of trials in which they first used their right index finger to perform center-out target-directed movements. After these baseline trials, we then applied a 45° clockwise (CW) rotation to the viewed cursor, allowing investigation of learning with the right hand. Following this, we briefly assessed participants' re-aiming strategy associated with their learning.

Each trial started with the participants moving the cursor (3 mm radius cyan circle) into the start position (4 mm radius white circle) in the center of the screen by sliding their index finger on the tablet. To guide the cursor to the start position, a ring centered around the start position indicated the distance between the cursor and the start position. The cursor became visible when its center was within 8 mm of the center of the start position. After the cursor was held within the start position for 0.5 s, a target (5 mm radius red circle) was shown on top of a grey ring with a radius of 60 mm (i.e., the target distance) centered around the start position. The target was presented at one of eight locations, separated by 45° (0, 45, 90, 135, 180, 225, 270 and

315°), in randomized bins of eight trials. Participants were instructed to hit the target with the cursor by making a fast finger movement on the tablet. They were instructed to 'slice' the cursor through the target to minimize online corrections during reach. If the movement was initiated (i.e., the cursor had moved fully out of the start circle) before the target appeared, the trial was aborted and a feedback text message ("too early") appeared centrally on the screen. In trials with correct timing, the cursor was visible during the movement to the ring and then became stationary for one second when it reached the ring, providing the participant with visual feedback of their endpoint reach error. If any part of the stationary cursor overlapped with any part of the target, the target turned green to indicate a hit. Each trial was terminated after 4.5 s, independent of whether the cursor had reached the target. After a delay of 1.5 s, allowing time to save the data, the next trial started with the presentation of the start position.

During the mock MRI training session (i.e., two weeks prior to the VMR MRI testing session), participants performed a practice block of 40 trials with their right hand with veridical feedback (i.e., no rotation was applied to the cursor). As with the RL task, this training session exposed participants to several key features of the task (e.g., use of the touchscreen tablet, trial timing, use of cursor feedback to correct for errors) and allowed us to establish adequate performance levels. At the beginning of the MRI testing session, but prior to the first scan being collected, participants re-acquainted themselves with the EL task by performing 80 practice trials with veridical cursor feedback (40 trials). Next, we collected an anatomical scan, followed by two fMRI experimental runs. The first run included a baseline block of 64 trials with veridical cursor feedback performed with the right hand, and the second run included a learning rotation block of 160 trials where cursor feedback was rotated clockwise by 45°.

##### *Behavioural data analysis*

**EL Task Processing.** Trials in which the reach was initiated before the target appeared (~4% of trials) or in which the cursor did not reach the target within the time limit (~5% of trials) were excluded from the offline analysis of hand movements. As insufficient pressure on the touchpad resulted in a default state in which the cursor was reported as lying in the top left corner of the screen, we excluded trials in which the cursor jumped to this position before reaching the target region (2% of trials). We then applied a conservative threshold on the movement and reaction times, removing the top 0.05% of trials across all subjects. As the EL task required the subject to determine the target location prior to responding, we also set a lower threshold of 100 ms on the reaction time. We then calculated the angle difference between the target position and the last sample before the cursor exceeded the target circle. This was then converted to an adaptation score by subtracting the angle of the rotation on each trial (0° during baseline, 45° during learning) to derive the actual aim direction on each trial.

**RL Task Processing.** Trials in which the cursor did not reach the target within the time limit were excluded from the offline analysis of hand movements (1% of trials). As insufficient pressure on the touchpad resulted in a default state in which the cursor was reported as lying in the top left corner of the screen, we excluded trials in which the cursor jumped to this position before reaching the target region (2% of trials). We then applied a threshold on the movement and reaction times, removing the top 0.05% of trials across all subjects. As the RL task did not involve response discrimination, we did not set a lower threshold on these variables.

**Learning Score Derivation (fPCA).** To derive a continuous measure capturing individual differences in overall learning proficiency across both EL and RL, we applied functional Principal Components Analysis (fPCA; 5) to participants' adaptation trajectories in the EL task, and scored trajectories in the RL task. As the RL task contained more trials than the EL task, we included the first and last 80 trials to ensure balanced representation of task trials in fPCA scores. This approach was chosen to preserve both early and late learning dynamics in the RL data, rather than simply truncating the final trials, which would have removed the late-learning phase most relevant to capturing individual differences in overall learning ability. This allowed us to identify the dominant modes of variation in how participants' performance evolved over the course of learning across both tasks, creating a behavioural index of generalized motor learning ability. For each participant, learning curves from both tasks were aligned along a common trial axis and concatenated to provide a unified representation of learning dynamics across both tasks. Learning trajectories were smoothed with a 5-trial median filter to reduce trial-level noise while preserving the overall shape of learning-related changes. Each participant's learning curve was then represented as a continuous function using a cubic B-spline basis (order 4), with smoothness enforced via a second-derivative penalty. This procedure converted discrete trial-wise performance into a smooth functional data object suitable for functional analysis.

We next performed fPCA on our set of participant-specific learning curves (6). Just as standard PCA estimates a low-dimensional subspace capturing variability in multivariate data, fPCA identifies a low-dimensional set of orthogonal functional components that capture dominant modes of variability across individuals (7). This approach is particularly well-suited for modelling learning, as it preserves the continuous temporal structure of performance while isolating shared patterns of inter-subject variability (for similar approaches see (8, 9). Our results yielded a set of functional principal components (fPCs), each explaining a decreasing proportion of the total variance in learning trajectories. fPC1 captured the majority of the variance (61.2%) in overall learning performance across tasks. Participants' loadings on this component were therefore used as a single continuous index of generalized learning ability, with higher scores reflecting more effective learning across combined tasks. All spline smoothing and fPCA procedures were implemented using the scikit-fda Python package (10).

As indicated above, we truncated the RL data to 160 trials to match with the EL data. However, to ensure that our results were not driven by this truncation, we also repeated fPCA using the full set of RL trials. Participants' fPC1 scores derived from the truncated and full RL datasets were significantly correlated, confirming that our dimensionality reduction approach was robust to this preprocessing choice (see Supplementary Figure 1A). In addition, to assess the extent to which each task contributed to our generalized learning measure, we performed fPCA separately on EL and RL learning trajectories. Here, participants' task-specific fPC1 scores were then correlated with the combined-task fPC1 scores, revealing strong associations for both tasks. This indicates that the shared fPC1 captures a common axis of learning variability expressed across both EL and RL (see Supplementary Fig. 2)

##### *MRI Acquisition*

Participants were scanned using a 3-Tesla Siemens TIM MAGNETOM Trio MRI scanner located at the Centre for Neuroscience Studies, Queen's University (Kingston, Ontario, Canada). Functional MRI volumes were acquired using a 32-channel head coil and a T2*-weighted single-shot gradient-echo echo-planar imaging (EPI) acquisition sequence (time to repetition (TR) = 2000 ms, slice thickness = 4 mm, in-plane resolution = 3 mm x 3 mm, time to echo (TE) = 30 ms, field of view = 240 mm x 240 mm, matrix size = 80 x 80, flip angle = 90°, and acceleration factor (integrated parallel acquisition technologies, iPAT) = 2 with generalized auto-calibrating partially parallel acquisitions (GRAPPA) reconstruction). Each volume comprised 34 contiguous (no gap) oblique slices acquired at a 30° caudal tilt with respect to the plane of the anterior and posterior commissure (AC-PC), providing whole-brain coverage of the cerebrum and cerebellum. Each of the task-related scans included an additional 8 imaging volumes at both the beginning and end of the scan. On average, each of the MRI testing sessions lasted approximately 2 hours.

At the beginning of the RL MRI testing session, a T1-weighted ADNI MPRAGE anatomical was also collected (TR = 1760 ms, TE = 2.98 ms, field of view = 192 mm x 240 mm x 256 mm, matrix size = 192 x 240 x 256, flip angle = 9°, 1 mm isotropic voxels). This was followed by a series of Diffusion-Weighted scans, wherein we acquired two sets of whole-brain diffusion-weighted volumes (30 directions, b = 1000 s $mm^{-2}$, 65 slices, voxel size = 2 x 2 x 2 $mm^3$, TR = 9.3 s, TE = 94 ms) plus 2 volumes without diffusion-weighting (b = 0 s $mm^{-2}$). Next, we collected a resting-state scan, wherein 300 imaging volumes were acquired. For the baseline and learning scans during RL testing, 222 and 612 imaging volumes were acquired, respectively.

At the beginning of the EL MRI testing session, we gathered high-resolution whole-brain T1-weighted (T1w) and T2-weighted (T2w) anatomical images (in-plane resolution 0.7 x 0.7 $mm^2$; 320 x 320 matrix; slice thickness: 0.7 mm; 256 AC-PC transverse slices; anterior-to-posterior encoding; 2x acceleration factor; T1w TR 2400 ms; TE 2.13 ms; flip angle 8°; echo spacing 6.5 ms; T2w TR 3200 ms; TE 567 ms; variable flip angle; echo spacing 3.74 ms). These protocols were selected on the basis of protocol optimizations designed by (11). For the baseline and learning functional scans, 204 and 492 imaging volumes were acquired, respectively. An additional 252 imaging volumes were collected during a transfer phase that is not analyzed in the current study.

##### *fMRI Preprocessing*

Results included in this manuscript come from preprocessing performed using fMRIPrep 20.1.1 (12, 13); RRID:SCR_016216), which is based on Nipype 1.5.0 (14, 15; RRID:SCR_002502).

**Anatomical data preprocessing.** A total of 2 T1-weighted (T1w) images were found within the input BIDS dataset. All of them were corrected for intensity non-uniformity (INU) with N4BiasFieldCorrection (16), distributed with ANTs 2.2.0 (17, RRID:SCR_004757). The T1w-reference was then skull-stripped with a Nipype implementation of the antsBrainExtraction.sh workflow (from ANTs), using OASIS30ANTs as target template. Brain tissue segmentation of cerebrospinal fluid (CSF), white-matter (WM) and gray-matter (GM) was performed on the brain-extracted T1w using fast (FSL 5.0.9, RRID:SCR_002823, 18). A T1w-reference map was computed after registration of 2 T1w images (after INU-correction) using mri_robust_template (FreeSurfer 6.0.1, 19). Brain surfaces were reconstructed using recon-all (FreeSurfer 6.0.1, RRID:SCR_001847, 20), and the brain mask estimated previously was refined with a custom variation of the method to reconcile ANTs-derived and FreeSurfer-derived segmentations of the cortical gray-matter of Mindboggle (RRID:SCR_002438, 21). Volume-based spatial normalization to two standard spaces (MNI152NLin6Asym, MNI152NLin2009cAsym) was performed through nonlinear registration with antsRegistration (ANTs 2.2.0), using brain-extracted versions of both T1w reference and the T1w template. The following templates were selected for spatial normalization: FSL's MNI ICBM 152 non-linear 6th Generation Asymmetric Average Brain Stereotaxic Registration Model (22, RRID:SCR_002823; TemplateFlow ID: MNI152NLin6Asym), ICBM 152 Nonlinear Asymmetrical template version 2009c (23, RRID:SCR_008796; TemplateFlow ID: MNI152NLin2009cAsym).

**Functional data preprocessing.** For each of the 10 BOLD runs found per subject (across all tasks and sessions), the following preprocessing was performed. First, a reference volume and its skull-stripped version were generated using a custom methodology of fMRIPrep. Head-motion parameters with respect to the BOLD reference (transformation matrices, and six corresponding rotation and translation parameters) were estimated before any spatiotemporal filtering using mcflirt (FSL 5.0.9, 24). BOLD runs were slice-time corrected using 3dTshift from

AFNI 20160207; 25, RRID:SCR_005927). Susceptibility distortion correction (SDC) was omitted. The BOLD reference was then co-registered to the T1w reference using bbregister (FreeSurfer) which implements boundary-based registration (26). Co-registration was configured with six degrees of freedom. The BOLD time-series were resampled onto the following surfaces (FreeSurfer reconstruction nomenclature): fsaverage5, fsaverage. The BOLD time-series (including slice-timing correction when applied) were resampled onto their original, native space by applying the transforms to correct for head-motion. These resampled BOLD time-series will be referred to as preprocessed BOLD in original space, or just preprocessed BOLD. The BOLD time-series were resampled into standard space, generating a preprocessed BOLD run in MNI152NLin6Asym space. Grayordinates files (27) containing 91k samples were also generated using the highest-resolution fsaverage as intermediate standardized surface space. Automatic removal of motion artifacts using independent component analysis (ICA-AROMA, 28) was performed on the preprocessed BOLD on MNI space time-series after removal of non-steady state volumes and spatial smoothing with an isotropic, Gaussian kernel of 6 mm FWHM (full-width half-maximum). Corresponding "non-aggressively" denoised runs were produced after such smoothing. Additionally, the "aggressive" noise-regressors were collected and placed in the corresponding confounds file. Several confounding time-series were calculated based on the preprocessed BOLD: framewise displacement (FD), DVARS and three region-wise global signals. FD was computed using two formulations following Power (absolute sum of relative motions, 29) and Jenkinson (relative root mean square displacement between affines, 24). FD and DVARS were calculated for each functional run, both using their implementations in Nipype (following the definitions by 29). The three global signals were extracted within the CSF, the WM, and the whole-brain masks. Additionally, a set of physiological regressors were extracted to allow for component-based noise correction (CompCor, 30). Principal components were estimated after high-pass filtering the preprocessed BOLD time-series (using a discrete cosine filter with 128 s cut-off) for the two CompCor variants: temporal (tCompCor) and anatomical (aCompCor). tCompCor components were then calculated from the top 5% variable voxels within a mask covering the subcortical regions. This subcortical mask was obtained by heavily eroding the brain mask, which ensures it does not include cortical GM regions. For aCompCor, components were calculated within the intersection of the aforementioned mask and the union of CSF and WM masks calculated in T1w space, after their projection to the native space of each functional run (using the inverse BOLD-to-T1w transformation). Components were also calculated separately within the WM and CSF masks. For each CompCor decomposition, the k components with the largest singular values were retained, such that the retained components' time series were sufficient to explain 50 percent of variance across the nuisance mask (CSF, WM, combined, or temporal). The remaining components were dropped from consideration. The head-motion estimates calculated in the correction step were also placed within the corresponding confounds file. The confound time series derived from head motion estimates and global signals were expanded with the inclusion of temporal derivatives and quadratic terms for each (31). Frames that exceeded a threshold of 0.5 mm FD or 1.5 standardised DVARS were annotated as motion outliers. All resamplings were performed with a single interpolation step by composing all the pertinent transformations (i.e. head-motion transform matrices, susceptibility distortion correction when available, and co-registrations to anatomical and output spaces). Gridded (volumetric) resamplings were performed using antsApplyTransforms (ANTs), configured with Lanczos interpolation to minimize the smoothing effects of other kernels (Lanczos 1964). Non-gridded (surface) resamplings were performed using mri_vol2surf (FreeSurfer).

**Regional timeseries extraction.** For each task, participant, and functional run, the average BOLD timeseries data was computed from the grayordinate timeseries for (1) each of the 400 cortical regions defined by the Schaefer 400-parcellation (32), (2) each of the 32 subcortical regions predefined according to the Tian scale II, 3T subcortical atlas (33), and (3) 32 cerebellar regions predefined by the Nettekoven 32-region cerebellar atlas (34). Region timeseries data were denoised using the above-mentioned confound regressors in conjunction with the discrete cosine regressors (128 s cut-off for high-pass filtering) produced from fMRIPrep and low-pass filtering using a Butterworth filter (100 s cut-off) implemented in Nilearn. Finally, all region timeseries were z-scored. See our previous work (9, 35) for similar approaches.

##### *Neuroimaging data analysis*

**Covariance estimation and centering.** For each participant, regional timeseries data from task-based functional scans were segmented into six equal-length task epochs (Base, Early, and

Late for both EL and RL tasks; 198 imaging volumes per epoch) after discarding the first six volumes to avoid scanner equilibrium effects. In addition, full resting-state scans were retained as a baseline reference for intrinsic functional architecture (297 imaging volumes). For each participant and task epoch, we estimated FC matrices by computing region-wise covariance matrices using the Ledoit-Wolf estimator (36). All task epochs were matched in length to ensure that covariance estimates were not biased by differences in timeseries duration.

To isolate learning-related changes while minimizing subject-specific differences in FC, we centered task-based covariance matrices using the Riemannian framework described by (37), which leverages the natural geometry of the space of covariance matrices (8, 38). In brief, this process involved adjusting the covariance matrices of each participant to have a common mean, equivalent to the overall mean covariance, thus removing subject-specific variations in FC. First, a grand mean covariance matrix, S_gm, was computed by taking the geometric mean covariance matrix across all participants and epochs. Then, for each participant i we computed the geometric mean covariance matrix across task epochs, S_i, and each task epoch covariance matrix S_ij was projected onto the tangent space at this mean participant covariance matrix S_i to obtain a tangent vector

$$T_{ij} = \overline{S}_i^{1/2} log(\overline{S}_i^{-1/2} S_{ij} \overline{S}_i^{-1/2}) \overline{S}_i^{1/2},$$

where log denotes the matrix logarithm. We then transported each tangent vector to the grand mean using the transport proposed by (37), obtaining a centered tangent vector

$$T_{ij}^c = G T_{ij} G^T,$$

where $G = \overline{S}_{gm}^{1/2} \overline{S}_i^{-1/2}$. Finally, we projected each centered tangent vector back onto the space of covariance matrices, to obtain the centered covariance matrix

$$S_{ij}^c = \overline{S}_{gm}^{1/2} exp(\overline{S}_{gm}^{-1/2} T_{ij}^c \overline{S}_{gm}^{-1/2}) \overline{S}_{gm}^{1/2},$$

where *exp* denotes the matrix exponential. For the benefits and general necessity of this centering approach, see (38). See also our previous work (9, 35, 39) for similar approaches.

**Manifold construction.** Following the estimation and centering of epoch- and task-specific covariance matrices, we applied manifold learning techniques to characterize large-scale network reconfiguration (35, 40–42). This approach embeds the connectivity profile of all brain regions into a low-dimensional space, providing data-driven, global views of distributed network dynamics without requiring predefined seed regions or a priori network models. As such, it provides complementary insights to methods focused on univariate activation or specific task-modulated pairwise interactions (e.g., Psychophysiological Interaction (PPI), Dynamic Causal Modeling (DCM); 43, 44).

Connectivity manifolds were constructed for each centered covariance matrix using the following procedure. First, consistent with prior work (35, 40–42), we applied row-wise thresholding to retain the top 10% of connections in each row. Cosine similarity was then computed between all pairs of rows to generate an affinity matrix, reflecting the similarity between regional connectivity profiles. Next, we performed principal components analysis (PCA) to obtain a set of principal components (PCs) that provide a low-dimensional representation of the connectivity structure (i.e., connectivity gradients). We selected PCA as our dimensionality reduction technique based on recent research indicating that PCA offers improved reliability over non-linear dimensionality reduction methods (40).

To provide a basis for comparing changes in functional network architecture across tasks and learning states, we constructed a template manifold from participants' resting-state connectivity data. This common resting-state matrix was derived by first computing, within each subject, a covariance matrix, and then computing the geometric mean across these resting-state matrices. We aligned all individual manifolds (37 participants x 6 epochs; 222 total) to this common baseline template manifold using Procrustes alignment. All analyses on the aligned manifolds were performed using the top five PCs, which cumulatively explained 65.4% of the total variance in the template manifold. Notably though, our observed effects are robust to changes in the number of PCs selected (see Supplementary Fig. 8). This approach enabled us to examine both task-specific and domain-general learning-related changes in manifold structure. See our previous work for a description of similar approaches (9, 35, 39, 45).

**Manifold eccentricity.** Recent work has quantified the embedding of brain regions in low-dimensional connectivity spaces using Euclidean distance metrics (9, 35, 46, 47). In the current work, we calculated 'manifold eccentricity' as the Euclidean distance between each region's coordinate in the aligned five-dimensional manifold space and the manifold centroid (defined as the origin: [0,0,0,0,0]; 45). This measure provides a scalar index quantifying how typical or atypical a region's whole-brain FC profile is relative to the average profile represented by the centroid. Regions with highly distinct connectivity patterns, often reflecting functional specialization, tend to lie further from the centroid, resulting in higher eccentricity, while regions sharing connectivity features across multiple systems lie closer, yielding lower eccentricity. In network neuroscience terms, higher eccentricity reflects greater functional segregation (i.e., a more distinct connectivity profile, often characterized by within-system coupling), whereas lower eccentricity reflects greater functional integration (i.e., a profile more similar to the whole-brain average, consistent with broader cross-system coupling; 45, 46). Eccentricity captures the relative distinctiveness of a region's connectivity pattern within the manifold and does not simply reflect overall connectivity strength or magnitude.

To empirically validate this interpretation within our data, we examined the relationship between baseline, resting-state eccentricity and established graph-theoretical measures of network topology (see Supplementary Fig. 4). These measures were computed on the row-wise thresholded template connectivity matrix and included node strength (the sum of connectivity weights), within-module degree z-score (relative within-network centrality), and participation coefficient (distribution of connectivity across networks; 48). Eccentricity showed significant associations with these measures, supporting its interpretation as an index of segregation versus integration. For our main analyses, we computed manifold eccentricity for each brain region, participant, and task epoch, allowing us to statistically test for manifold expansions (increases in eccentricity) and contractions (decreases in eccentricity) throughout Base, Early, and Late learning, as well as across tasks. See our previous work (9, 35, 39, 45) for similar approaches. While our primary analyses were conducted using the top five PCs, the inclusion of additional components did not meaningfully affect our results (see Supplementary Fig. 8).

**Eccentricity analysis.** To examine changes in manifold eccentricity across epochs, we conducted a 2 (Task: EL, RL) x 3 (Epoch: Base, Early, Late) repeated measures ANOVA (rmANOVA) separately for each brain region. This allowed us to test for main effects of Task, main effects of Epoch, and their interactions on regional eccentricity. To account for multiple comparisons across regions and effects, p-values were corrected using a false-discovery rate (FDR) procedure ($q < 0.05$; 49). For regions showing a significant main effect or interaction, we conducted follow-up paired samples t-tests to characterize differences between specific tasks and/or epochs. See our previous work for similar implementations of this analytic framework (9, 35, 39, 45). To determine whether our observed effects in manifold eccentricity could be explained by changes in univariate activation, we quantified the mean BOLD signal within regions showing significant main effects of Task and Epoch. This analysis revealed that patterns of mean BOLD activation did not mirror the connectivity-driven effects observed in manifold eccentricity, suggesting that our findings are not reducible to simple changes in regional activation magnitude (see Supplementary Fig 9).

**Investigating seed connectivity across the PCC.** While most regions in our eccentricity analyses exhibited a main effect of either Task or Epoch, six posterior cingulate cortex (PCC) regions showed significant main effects of both. To better understand the underlying covariance changes driving these reconfigurations in manifold eccentricity, we performed a series of seed-based connectivity contrasts across Tasks and Epochs separately. For the six PCC regions, we first computed averaged eccentricity values for each Task and Epoch to characterize the direction and magnitude of the observed main effects. We then generated subject-level seed-based connectivity maps, aggregated across all PCC regions for (1) Task-effects, collapsing across epochs (EL vs RL), and (2) Epoch-effects, collapsed across tasks (Early > Base, Late > Early, Late > Base). For each contrast, we used paired-samples t-tests across subjects to quantify differences in connectivity patterns. As manifold eccentricity reflects a multivariate shift in whole-brain connectivity structure — meaning that no single pairwise connection constitutes a direct test of what drives a manifold change — we visualized the resulting unthresholded t-maps to capture the full spatial pattern of connectivity changes underlying eccentricity differences.

To further summarize the effects at the whole-brain network level, we constructed polar plots for the PCC seed regions by averaging t-values across regions according to their cortical network assignments (32), the subcortex (33) and the cerebellum (34).

**Correlating whole-brain eccentricity changes with overall learning performance.** To determine whether learning-related changes in whole-brain functional architecture were associated with individual differences in general learning ability, we related changes in regional manifold eccentricity to participants' overall behavioural learning scores (fPC1). For each brain region, we computed within-subject eccentricity change scores for each epoch contrast (Early > Base, Late > Early, Late > Base). We then quantified the relationship between these regional change scores and fPC1 across participants using Pearson correlations, separately for each contrast.

As ROI-wise cortical effects are often spatially autocorrelated, we additionally tested these associations at the network-level (32) using spatial null models. Cortical parcel-wise values were aggregated within the Yeo 7-network parcellation (32) to obtain an empirical mean correlation for each network and contrast. Statistical significance was assessed using spin permutation testing (1,000 rotations), which preserves large-scale spatial autocorrelation structure. For each network, a two-sided spin-based p-value was computed by comparing the empirical network mean to its null distribution. In parallel, we performed one-sample t-tests to determine whether parcel-wise correlations within each network differed significantly from zero. Networks were considered significant if they met both criteria (spin-based $p < 0.05$ and zero-test $p < 0.05$). As spin-based null models are defined only for surface-based cortical data, cerebellar and subcortical regions were excluded from this analysis.

To further illustrate significant network-level findings, we selected one cortical ROI per network and contrast based on the strongest correlation within that network. For the Limbic network, which showed significant effects in two epoch contrasts, we identified a single ROI exhibiting a strong positive association from Base to Early learning, and a strong negative association from Early to Late learning, allowing for a full visualization of the state-dependent reversals within the same region. For each selected ROI, participant-level scatterplots were generated to depict the relationship between manifold eccentricity and overall learning ability.

**Seed connectivity analysis.** To understand the underlying learning-related changes in regional covariance that give rise to the observed changes in manifold eccentricity, we conducted targeted seed-based connectivity analyses using the representative regions identified from the network-level effects described above. For each selected seed region and participant, we derived seed-based connectivity maps for Base, Early, and Late epochs. Paired samples t-tests were performed across participants to contrast connectivity between epoch comparisons (Early > Base, Late > Early, Late > Base). Consistent with our interpretation of eccentricity as a multivariate measure of connectivity, we visualized unthresholded whole-brain t-maps to depict the full spatial pattern of connectivity differences underlying observed eccentricity changes.

To further illustrate these effects, we generated participant-level scatterplots depicting eccentricity changes across epochs for each seed. We also constructed polar plots summarizing connectivity differences at the network-level by averaging t-values according to their cortical network assignments (32), the subcortex (33) and the cerebellum (34). These analyses were intended to provide qualitative insight into the distributed connectivity changes of representative regions showing significant eccentricity effects (for similar visualization-based approaches see (9, 35, 45).

**Relating seed connectivity to overall learning performance.** To provide a descriptive visualization of how learning-related behavioural differences mapped onto large-scale FC changes, we related seed-based connectivity changes to participants' overall learning scores (fPC1 scores). This analysis was intended to support qualitative interpretation and complement our primary manifold eccentricity findings, rather than serve as a formal inferential test.

For each seed and task epoch, connectivity values were averaged within the Yeo 7-network parcellation (32), as well as within the subcortex (33) and cerebellum (34). For each participant, connectivity change scores were computed along three epoch contrasts (Early > Base, Late > Early, Late > Base). For each seed region, we quantified the association between connectivity changes and behavioural learning performance by computing Pearson correlations across participants between networks. Results were displayed both in surface space and using polar plots.


##### *Funding Information*

This work was supported by operating grants from the Canadian Institutes of Health Research Grant (PJT175012) and the Natural Sciences and Engineering Research Council (RGPIN-2017-04684; RGPIN-2024-06773), awarded to J.P.G. The funders had no role in study design, data collection and analysis, decision to publish, or preparation of the manuscript.

##### *Acknowledgments*

The authors would like to thank Martin York and Sean Hickman for technical assistance, and Don O'Brien for assistance with data collection.
